# Family-based association analysis identifies variance-controlling loci without confounding by genotype-environment correlations

**DOI:** 10.1101/175596

**Authors:** Dalton Conley, Rebecca Johnson, Ben Domingue, Christopher Dawes, Jason Boardman, Mark Siegal

## Abstract

The propensity of a trait to vary within a population may have evolutionary, ecological, or clinical significance. In the present study we deploy sibling models to offer a novel and unbiased way to ascertain loci associated with the extent to which phenotypes vary (variance-controlling quantitative trait loci, or vQTLs). Previous methods for vQTL-mapping either exclude genetically related individuals or treat genetic relatedness among individuals as a complicating factor addressed by adjusting estimates for non-independence in phenotypes. The present method uses genetic relatedness as a tool to obtain unbiased estimates of variance effects rather than as a nuisance. The family-based approach, which utilizes random variation between siblings in minor allele counts at a locus, also allows controls for parental genotype, mean effects, and non-linear (dominance) effects that may spuriously appear to generate variation.

Simulations show that the approach performs equally well as two existing methods (squared Z-score and DGLM) in controlling type I error rates when there is no unobserved confounding, and performs significantly better than these methods in the presence of confounding. Using height and BMI as empirical applications, we investigate SNPs that alter within-family variation in height and BMI, as well as pathways that appear to be enriched. One significant SNP for BMI variability, in the MAST4 gene, replicated. Pathway analysis revealed one gene set, encoding members of several signaling pathways related to gap junction function, which appears significantly enriched for associations with within-family height variation in both datasets (while not enriched in analysis of mean levels). We recommend approximating laboratory random assignment of genotype using family data and more careful attention to the possible conflation of mean and variance effects.

## Introduction

### From effects of loci on trait means to effects of loci on trait variance

The extent to which a complex trait varies in a population is a product of mutation, genetic drift and natural selection, as well as environmental variation and its interaction with genotype. Moreover, genotypes may differ in their propensities to vary. That is, particular genotypes might be more sensitive than others to changes in the environment or to the effects of new mutations. Sensitivity to the environment could come in the form of phenotypic plasticity, whereby stereotyped phenotypic changes occur under particular circumstances, or in the form of developmental instability, whereby random fluctuations in the internal or external environment lead to different phenotypic outcomes ( [23,29,52]).

Sensitivity to the effects of mutations is related to cryptic genetic variation. In a population in which individuals are relatively insensitive to the effects of mutations, allelic variation may accumulate, while not presenting phenotypic effects. Replacing an allele that confers less sensitivity to mutational effects with one that confers more sensitivity (or introducing an environmental perturbation with similar effect) will cause the accumulated variation to have phenotypic consequences ( [33,54]). This release of cryptic genetic variation might have major implications for the adaptation of populations to environmental change, as well as for the genetics of human complex traits ( [9,32,34,81]). Indeed, it has been proposed that release of cryptic genetic variation might be responsible for the increased prevalence of human “diseases of modernity” such as diabetes ( [32]).

Model organisms have been used to identify genes that control sensitivity to mutations or the environment [54]; [71]). Two approaches have been taken. One approach is to test for differences in phenotypic variance between wild-type and mutant strains. For example, the molecular chaperone Hsp90, when impaired, reveals cryptic genetic variation in the fly Drosophila melanogaster, the flowering plant Arabidopsis thaliana, the fish Danio rerio and the budding yeast Saccharomyces cerevisiae ( [45,61,63,84]).

Evidence on whether Hsp90 controls environmental sensitivity is mixed ( [39,66,70]) However, screens of the S. cerevisiae genome identified hundreds of genes that, when mutated, cause increased variation in cell morphology among isogenic cells raised in the same environment. That is, these mutations increased sensitivity to fluctuations in the internal or external environment ( [8, 51]). A test of one of these genes, which showed a major increase in environmental sensitivity upon deletion, revealed a high degree of epistasis with new mutations but no net increase in mutational sensitivity upon deletion ( [52,62,62]). A similar result was found for impairment of Hsp90 ( [30]). In general, the relationship between suppression of the effects of environmental variation and suppression of the effects of mutational variation is unclear ( [54,71]).

The second approach to identifying variance-controlling genes is to use linkage or association analysis to map natural genetic variants that confer differential sensitivity. Such searches for variance-controlling quantitative trait loci (vQTLs) have been conducted to identify determinants of microenvironmental sensitivity (i.e., inherent stochasticity or sensitivity to fluctuations within a nominally constant environment) and/or phenotypic plasticity (variation across controlled environments) of various phenotypes, including morphological and life-history traits as well as expression levels of individual genes, in a range of organisms including humans ( [5,7,26,27,35,40,44,46,59,65,69,74,77-79]). Natural genetic variation affecting sensitivity to other segregating alleles has been studied as well, and indeed there are natural variants of the Hsp90 gene that appear to reveal cryptic variation ( [17,68,72]).

### Existing methods for detecting vQTL

The first study claiming to map a locus that controls variance of a human trait examined body mass index (BMI) using data from 38 cohorts that participate in the GIANT consortium for genome-wide analysis (GWA) of single-nucleotide polymorphisms (SNPs) that affect human height ( [25])(although a prior study had examined variance in order to prioritize the search for gene-by-environment and gene-by-gene interaction effects [58]). Yang et al. computed a Z-score (inverse-normal transformation) for BMI for each of 133,154 individuals then performed GWA on the squared Z-score. This squared Z-score captures the magnitude of each individual’s deviation from the mean phenotype and therefore is meant to act as an individual-based measure of variance. In this discovery sample they found SNPs in the FTO gene and the RCOR1 gene that appear to control variation in human BMI. In a replication sample of 36,327 individuals from 13 cohorts, one SNP in the FTO gene was confirmed [25].

### Challenge one: isolating loci that affect variance in a trait from loci that affect mean of a trait

There are several challenges with association analyses that identify vQTL. The first issue is mean-variance confounding. A given locus can have: 1) effects on mean levels of a trait (mean effects), 2) effects on variance in a trait (variance effects), or 3) effects on both mean levels of a trait and variance in a trait. For isolating variance effects, it is critical to check that any detected effects on “variance” adequately distinguish between the three types of loci. In particular, this conflation of variance and mean effects was a concern in the analysis of Yang et al. ( [25]) because normalized variation scores will tend to be higher for populations with higher mean levels and because FTO is one of the most well-established genes that affects the mean level of BMI in human populations ( [4,14,16,22,28,37,41,67,80]). Research examining MMP3 protein levels in cerebrospinal fluid found similar overlap, where SNPs in linkage disequilibrium with a locus well-established in predicting mean levels of the trait (rs679620 of the MMP3 gene) were associated with both higher mean levels and higher variance in the trait ( [12]).

Yang et al. addressed the effect of mean BMI by showing that there is no global correlation between mean effects of SNPs and their variance effects. However, it must be considered that a lack of correlation between mean and variance effects across the genome could be caused merely by the fact that the vast majority of SNPs have negligible or nonexistent effects on both mean and variation. In the present study, we first show through simulation that the squared Z-score method has an inflated type I error rate and detects variance effects for a trait simulated to have mean effects only. Then, we show through an empirical analysis of the top-scoring SNPs from the Yang study that significant confounding between mean and variance effects does in fact exist.

In addition to mean-variance confounding, methods to investigate vQTL must address several other issues, some of which are shared with the estimation of mean effects and others of which, such as mean-variance confounding, are unique to or particularly acute in the case of variance effects ( [65]). Methods for vQTL analysis beyond the squared Z-score method can be grouped into: 1) non-parametric methods that test whether the three genotypes at a biallelic locus (minor-allele homozygote, heterozygote, and major-allele homozygote) have equal or unequal variance, and 2) parametric methods that relax the assumption in GWA linear regressions that the residual error is identically distributed across all genotypes.

Non-parametric methods include Levene’s test, which uses a test statistic derived from the squared difference between an individual’s level of a trait and the genotype-level mean or median, the latter of which is used to make the method more robust to a non-normally distributed trait ( [58,75]) and the Fligner-Killeen (FK) test, which is similar to Levene’s test, in that it uses the absolute difference between an individual’s level of a trait and the genotype-level median, but then computes the test statistic based on ranks of these differences. The FK test can be used either as a standalone test for variance effects ( [27]), or as the test statistic for the scale component of the Lepage or other joint scale-location test ( [73]). More recently proposed non-parametric tests consider not only the variance of the trait distribution but other features as well, such as skew ( [6,38]).

The main drawback of non-parametric tests for detecting vQTL’s is that the tests, which compute differences between discrete genotype groups, cannot directly control for important covariates, such as age, sex, and population structure. Some adopt a two-stage regression procedure for including covariate controls: first the trait is regressed on covariates, then the variance test of interest is performed on the residualized dependent variable ( [38,83]). However, two-stage procedures have been shown in simulations to reduce power and induce bias ( [12]).

### Challenge two: controlling for unobserved confounders that can bias the estimate on minor allele count

This problem-how to include covariates in a way that does not induce bias-is acute because of the importance of two types of controls when investigating variance-controlling loci: control for population stratification and controls for nonrandom association between genotype and environment. The former control is needed in all GWA analyses to separate the effect of any particular locus from the effects of all other loci shared by virtue of common ancestry. The latter control might be particularly relevant to vQTL analyses because mean effects of genotypes might impact the environment that is experienced, which might in turn impact variance ( [13]).

Parametric approaches that use generalized linear models to jointly estimate the mean and variance of a trait address this problem ( [11,12,64]); these models can include controls for population stratification as well as controls for observed covariates that influence genetic distribution into variance-affecting environments. The double generalized linear model (DGLM) approach begins with the typical linear model for estimating mean effects where the residual variance is the same across genotypes, then the model is relaxed to allow residual variance to differ by genotype and to incorporate non-genetic covariates that might contribute to residual variance; it iterates between estimating parameters for the mean versus parameters for the variance until convergence ( [64]). DGLM thus allows joint estimation of mean effects and variance effects, attempting to address mean-variance confounding, and permits controls for population stratification directly in the model. The main drawback of DGLM is that, similar to other methods that control for confounding by controlling for *observed covariates* (age; sex; the top principal components) correlated with both the individual’s genotype and the outcome variable, the method cannot control for unobserved confounding that may bias the estimate of a SNP’s effect.

In particular, two types of unobserved confounders may be correlated both with an individual’s genotype and mean or variance effects in a trait. First is population stratification. Population stratification is typically controlled for in these methods by inclusion of principal components of the sample population’s genotypes among the vectors of covariates predicting the mean and/or residual variance of a trait. This control is especially important for traits, such as BMI, that are expected to show considerable environment-dependence. Especially when pooling such traits across cohorts, there is a risk that systematic differences in environment correlate regionally with systematic differences in genetic variation. That is, it is plausible, due to population stratification, that any genetic signal is merely acting as a proxy for culture and environment-a potential confounder that has been well-illustrated by the “chopsticks gene” example ( [36]). Controlling for population stratification using principal components (PCs) addresses some confounding, but there is residual between-family confounding even with these controls. This residual confounding can occur when environmentally influential factors are not randomly distributed across families but also do not correlate with the eigenvectors in the genetic matrix.

The other critical covariate that might bias vQTL estimates is genotype-environment correlation (rGE). Genotype-environment correlations may be caused by niche construction, whereby individual organisms shape the environment (in a genotype-dependent way) ( [20,48,49,56,57])–dynamics we might expect to occur for phenotypes that have significant behavioral and environmental etiologies, such as BMI. As a result, genotypes may be associated with variance in BMI and other traits not through direct genetic effects but through interaction with alternative environments associated with variance such as more versus less sedentary lifestyles. An analogous situation that illustrates this potentially confounding genotype-by-environment interaction effect is that of caffeine consumption. A variant in a gene that encodes a caffeine-metabolizing enzyme can lead to greater variation with no effect on the mean through a mechanism of niche construction-i.e. individuals with the minor allele avoid coffee altogether, or if they are unable to do so, they end up drinking more than those with the major allele, thus leading to greater variance thanks to the coffee “environment” ( [13]).

Approaches such as DGLM can control for observed covariates that are correlated with genotype and influence construction of variance-affecting environments. However, these methods cannot control for unobserved differences that produce these correlations. The present paper uses a family-based model to control for these unobserved differences. Two existing family-based models allow for investigations of variance-based loci in samples among related as opposed to unrelated individuals, but do not leverage the family-based structure of the data to control for unobserved confounders that vary between families ( [11]; [82]). First is a family-based version of the likelihood ratio test that adds a random effect meant to capture familial correlation in a trait. Although the family-level random effect helps control for unobserved variation between families that may influence variance in a trait through pathways other than genotype, the model relies on the strong assumption of independence between these unobserved features of family and the observed covariates. We show via simulation (see Results) that when there is non-zero correlation between unobserved features of a family and observed covariates, random effects approaches generate biased estimates of a SNPs’ effects, confirming results shown in non-genetics contexts ( [21]). Similarly, DGLM in a sample containing monozygotic and dizygotic twins, which has the advantage of isolating non-genetic from genetic sources of variance, does not control for unobserved features that vary between families and that affect construction of variance-affecting environments ( [82]).

### Proposed solution: the sibling standard deviation method

We offer an alternative methodology-comparisons of variation within sibling sets while controlling for parental genotype-that does not assume independence between observed covariates and unobserved between-family differences. As a result, the method better approximates random assignment of genotype in a laboratory. Utilizing a regression-based framework, the approach retains the advantages of the DGLM and Bayesian regression approaches: the ability to include covariates and control for mean effects when estimating variance effects through estimation of parameters capturing both. The model uses sibling pairs as the unit of analysis and regresses the standard deviation of the sibling pair’s trait on the pair’s count of minor alleles with sibling pair-level controls that include controls for the mean level of the trait in the sibling pair, parental genotype, pair sex (MM or FM or FF), mean pair age, and the within-pair age difference (for the full model specification, as well as alternative specifications tested, see Methods). The control for the mean level of the trait in a sibling pair avoids inflated Type I error rates for SNPs that affect the mean of a trait but not the trait’s variance.

The proposed methodology, although restricted in applicability to datasets that have a family-based design with at least two offspring, makes a trade-off. The method has reduced statistical power because the sample size is halved when we treat siblings rather than individuals as the unit of analysis. However, the advantage is an estimate of a minor allele’s contribution to variance that, in the presence of unobserved confounders, correctly fails to find variance effects when a locus only has mean effects.

This is a particularly important trade-off to make when we wish to rule out gene-environment correlations across populations as well as population stratification as alternative explanations to variance-locus associations. As noted above, this is especially critical for a phenotype such as BMI and a locus such as FTO given that environment and behavior (such as sedentariness) alter the FTO-BMI relationship and may vary significantly across cohorts/societies ( [3]). In cases where the goal is not to study control of variance per se, but instead is to probe the existence of gene-environment or gene-gene interactions in a way that avoids the high-dimensional parameter space problems of traditional approaches, the use of vQTL approaches that have higher statistical power but also a higher rate of false positives for SNPs that affect the mean of a trait but not the variance might be warranted.

In addition to power, another important feature of an approach to detect vQTL is the flexibility to capture non-linear effects of alleles, which the DGLM, the parametric bootstrap-based likelihood ratio test, and Bayesian regressions allow for by allowing genotypes to be specified using three indicator variables to capture non-linearities. The present family-based approach is potentially susceptible to confounding of variance effects by non-linear effects of alleles, because the association mapping is done on the sibship unit so the genotype is represented as the total number of major or minor alleles of each sib pair. Therefore, if dominance were at play with respect to mean levels, then this might generate spurious effects on variance ( [76]). That is, if among heterozygotes there was no effect of an allele on mean levels but among homozygotes there was, then this itself would generate apparent, but spurious, effects on variance even when controlling for linear mean effects. However, by comparing subgroups among sibship pairs that have two minor alleles (out of four possible in total), we are able to check for this possibility. Specifically, by comparing those 2-minor allele pairs where both siblings are heterozygotes (i.e. each individual has one minor allele) with those where both are homozygotes (where one sibling has zero minor alleles and the other has two), we can rule out this possible statistical artifact of non-linear effects on mean levels. Although the possibility of non-linearities that do not reflect true variance effects can never be totally eliminated ( [76]), this approach guards against a primary form of non-linearity—dominance.

### Preview of the results

Below we report two sets of analyses. In simulations, we show how unobserved confounding can bias estimates of a SNP’s effect. Two approaches to estimating variance effects-the squared Z-score method and DGLM-display inflated type I error rates in the presence of this confounding. In contrast, the sibling standard deviation approach that we propose detects variance effects when these effects are present but, by controlling for the mean of the trait across siblings, correctly fails to find variance effects when only mean effects are present.

In an empirical application, we then use the sibling SD approach to perform genome-wide analyses of variance effects on two phenotypes: height and BMI. We replicate one genome-wide statistically suggestive hit for variation in BMI from the Framingham Heart Study (FHS) data, our discovery sample, in our replication sample, the Minnesota Twin Family Study (MTFS). We also test whether our potential variance-related alleles are merely reflecting dominance effects among heterozygotes; we find no evidence for this. Finally, we perform gene-based and pathway enrichment analysis. We find one pathway, related to gap junction function, that is significantly enriched in both our discovery and replication samples for associations with variance in height. We then discuss the implications of our findings for prior and future research.

## Results

### Simulations

The simulations are divided into two parts. First, we show that while we can address unobserved confounding when estimating the *mean* of a trait using a fixed effects estimator that identifies the effect of an allele off of between-sibling variation, combining this approach with two current approaches to variance detection-the squared Z-score method and DGLM-fails to correct for this bias. This introduces the challenge: how can we estimate the effect of an allele on trait variance in the presence of unobserved confounding? Part two evaluates the sibling SD method as a solution.

### Part one: two approaches to variance estimation in the absence versus presence of an unobserved confounder

When estimating mean effects, we can remove bias in the estimate of the effect of a minor allele caused by unobserved confounding by using a fixed effects estimator that demeans the outcome, genotype, and other observed covariates by the mean within a family ( [21]). Examining the trait simulated to have *neither mean nor variance effects*, S1 Fig and S1-S4 Tables show that in the presence of any unobserved confounding, other estimators (a random effects model that assumes zero correlation between unobserved confounders and observed covariates; a pooled regression model with one randomly sampled sibling) return biased estimates of the effect of an additional minor allele on the trait’s mean. More specifically, S1 Fig shows that across the 1000 replicates, we see upward bias in the random effects and pooled regression estimators when we move from the case of no confounding to the case of some confounding. S1 and S2 Tables, which present regression results for one randomly chosen replicate, show that the regressions estimate significant effects of an allele on the trait mean for the DV simulated to have no effects when confounding is present. S3 Table show that fixed effects correctly fail to find mean effects. S4 Table, which presents the results of a Hausman test comparing the null hypothesis that the estimated random effects coefficients are equal to the estimated fixed effects coefficients, rejects the null at both levels of confounding.

These results show how a fixed effects estimator that identifies the effect from sibling deviations from a family’s mean count of alleles recovers an unbiased estimate. How can we translate these findings from the case of investigating the effect of an additional minor allele on the trait’s mean (QTL) to investigating the effect of an additional minor allele on the trait’s variance (vQTL)?

One approach is to use existing methods for variance detection on a transformed version of the data. In particular, the fixed effects estimator is generated by demeaning the outcome, genotype, and other covariates by the mean across the grouping unit (family in this case), as represented as follows, where *i* indexes an individual, *j* indexes a family, *k* indexes a SNP, and *X* represents genotype and other covariates:

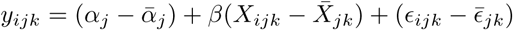

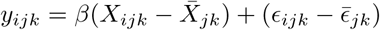

We can represent this demeaned version of the data more compactly as follows:

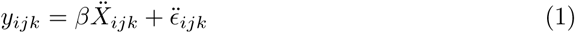

One approach to translating the FE estimator to the vQTL context is to use the transformed version of the data represented in Equation 1 with two existing methods for vQTL detection: the squared Z-score method ( [25]) and the DGLM ( [64]). We explore the properties of each approach in the sections that follow on two versions of the dependent variable:

1. *Dependent variable simulated to have mean effects only* : the dependent variable is simulated to have a positive relationship between the count of minor alleles and the mean of the outcome (but not the outcome’s variance) (see Methods)
2. *Dependent variable simulated to have variance effects only* : the dependent variable is simulated to have a positive relationship between the count of minor alleles and the variance of the outcome (but not the outcome’s mean) (see Methods)

### Squared Z-score results

S5 Table presents the results of regressing the squared Z-score (estimated separately by sex) on the minor allele count with controls for sex, age, and ancestry across the 1000 replicates. The table summarizes the percentage of replicates where the coefficient on the minor allele count ≠ 0 at *p* < 0.05. In the *absence* of an unobserved confounder, we see moderately inflated type I error rates (estimating variance effects when only mean effects are present in 7.1% of simulations without an ancestry control, 6.5% with an ancestry control). In the *presence* of an unobserved confounder, we see highly inflated type I error rates: 24.6% of simulations estimate variance effects when only mean effects are present in the model without an ancestry control, 24% with an ancestry control. S6 Table shows that this inflated type I error rate is only slightly lower when we estimate the model on demeaned data, falling to 20% of simulations that estimate *β* ≠ 0 for the trait simulated to have mean but not variance effects.

### DGLM results

The squared Z-score method generates one coefficient of interest that represents an allele’s contribution to variability in the form of increasing an individual’s squared Z-score. The DGLM method, which estimates a linear regression for the trait mean and a gamma regression from the squared residuals from the first model, generates two coefficients of interest: (*β* refers to coefficients on mean effects; *γ* refers to coefficients on variance effects). Table 1 summarizes the true *β* and *γ* for different types of simulated traits:

**Table 1.** Outcomes versus coefficients

| Simulated dependent variable | Coefficient on minor allele count from DGLM |
| --- | --- |
| Mean effects only | $\beta \neq 0; \gamma = 0$ |
| Variance effects only | $\beta = 0; \gamma \neq 0$ |

S7 Table looks at the case where the simulated DV has variance effects only (but no mean effects) across the 1000 replicates. The DGLM should estimate *β* = 0 for mean effects and *γ* ≠ 0 for variance effects. Therefore, the table summarizes the percentage of *β* ≠ 0 at the *p* < 0.05 level; if the model has a low type I error rate, this percentage should be low. The table also summarizes the percentage of *γ* ≠ 0 at the *p* < 0.05 level; if the model has low type II error rate, this percentage should be high.

The results show that in the presence of between-family confounding between the genotype and outcome variable, the non-transformed data incorrectly detects mean effects when the DV is simulated to only have variance effects (type I error). The model estimated on demeaned data has a lower false positive rate but still incorrectly detects mean effects (estimates *β* ≠ 0 at *p* < 0.05 for 20.4% of the simulations) when a trait is simulated to have variance effects only. More problematic, the model estimated on demeaned data fails to detect variance effects when these are present (type II error).

### Problems part one reveals

The previous section reveals a problem when choosing a method for detecting the effects of an additional minor allele on the mean or variance of a trait in the presence of unobserved confounding between an individual’s genotype and the trait. In particular, while the fixed effects estimator provides an unbiased estimate of the effect of minor alleles on the *mean* of a trait, de-meaning the data and then trying to estimate variance effects using the squared Z-score or DGLM approach leads to inflated rates of type I error when the loci has mean effects but no variance effects. In addition, the DGLM estimated on demeaned data fails to detect variance effects when these effects are present.

These problems point to the need for a method that corrects for bias caused by unobserved confounding but that is also able to detect effects of minor alleles on a trait’s variance when these effects are present. The next section investigates properties of the sibling standard deviation method as a proposed solution.

### Part two: properties of the sibling standard deviation method

To examine variance effects, we estimate the sibling standard deviation method (see Methods). We first summarize results from one randomly chosen replicate to highlight the important role played by controlling for the sibling mean of a trait, and then summarize results across the 1000 replicates.

### Results from one randomly chosen replicate

S8 Table, which does *not* control for parental genotype, and S9 Table, which *does* control for parental genotype, present two coefficients from the sibling SD model: the coefficient on the sibling minor allele count, which should significantly differ from zero for the traits simulated to have variance effects but should fail to differ from zero for the traits without these effects, and the coefficient on the sibling mean of the trait, which should differ from zero for the trait with mean effects. The results show that the sibling standard deviation method detects variance effects of a snp and properly rejects mean effects of a snp both in the absence and presence of family-level confounding. The results highlight that this power to reject false positives when a locus affects the mean and not the variance of a trait comes from controlling for the sibling mean of the trait, which is significant for the traits simulated to have mean effects.

Figure 1, which presents the sibling count of minor alleles and non-adjusted mean sibling standard deviation for pairs with that count for one randomly chosen replicate, highlights this pattern of an increase in the trait’s sibling standard deviation when variance effects are present and no increase in the trait’s sibling standard deviation in the presence of mean effects alone.

**Figure 1.**
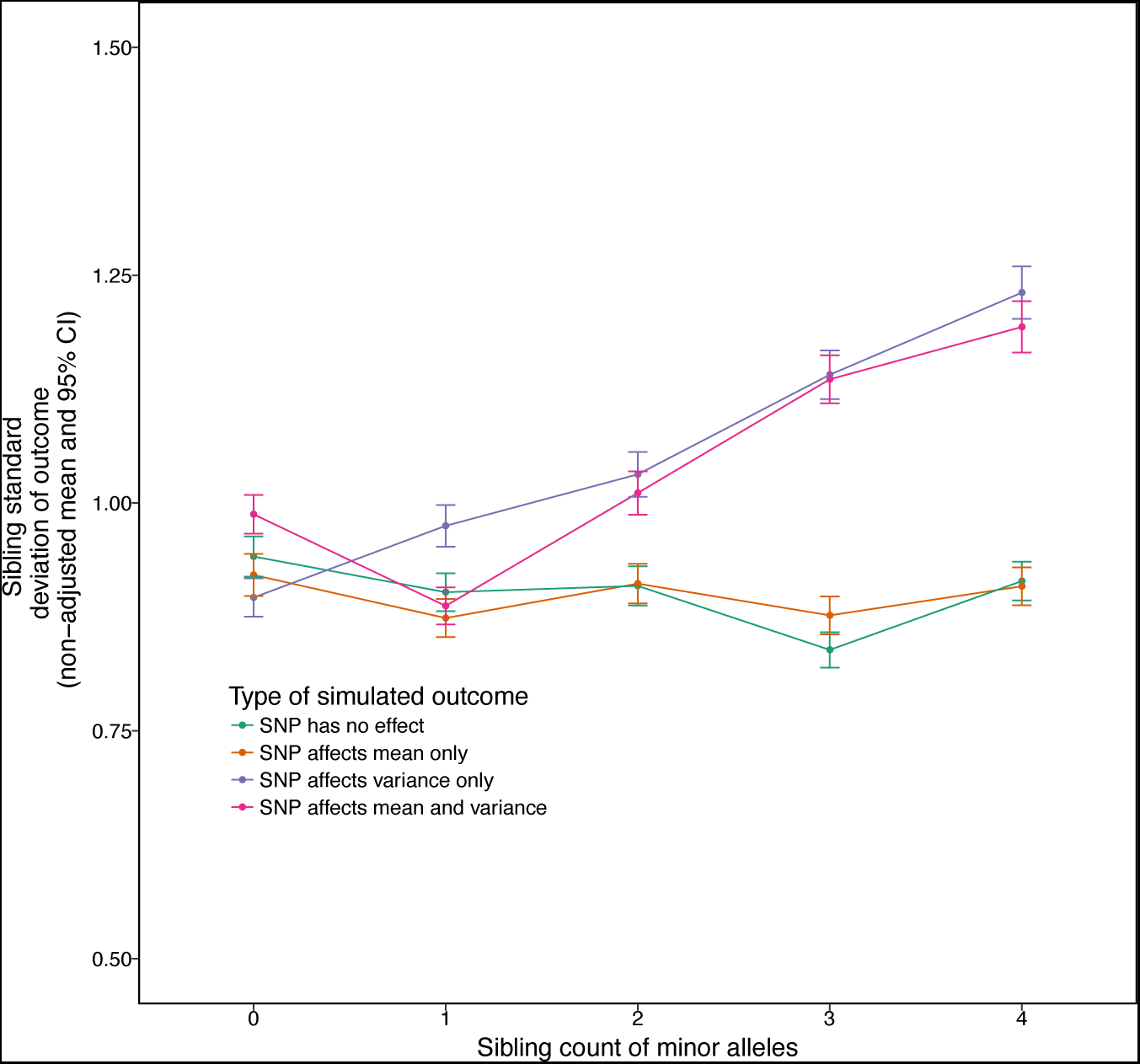
Raw means of sibling standard deviations (before age, sex, mean of trait, and parental genotype controls) by count of minor alleles. The graph shows that the sibling standard deviation increases with the count of minor allele snp’s that have effects on variance only, or effects on both the mean and variance of a trait, while stays flat for snp’s that only effect the mean or that not associated with the trait.

### Results across 1000 replicates

Figure 2 shows that the results from the previous method generalize across the replicates. The figure shows the distribution of *β* on sibling minor allele count from regressing the sibling standard deviation of a trait on this count for two traits: the trait simulated to have mean effects only and the trait simulated to have variance effects only. The figure shows the distribution is properly centered around zero for the trait with mean effects and properly centered around *β* ≠ 0 for the trait with variance effects.

**Figure 2.**
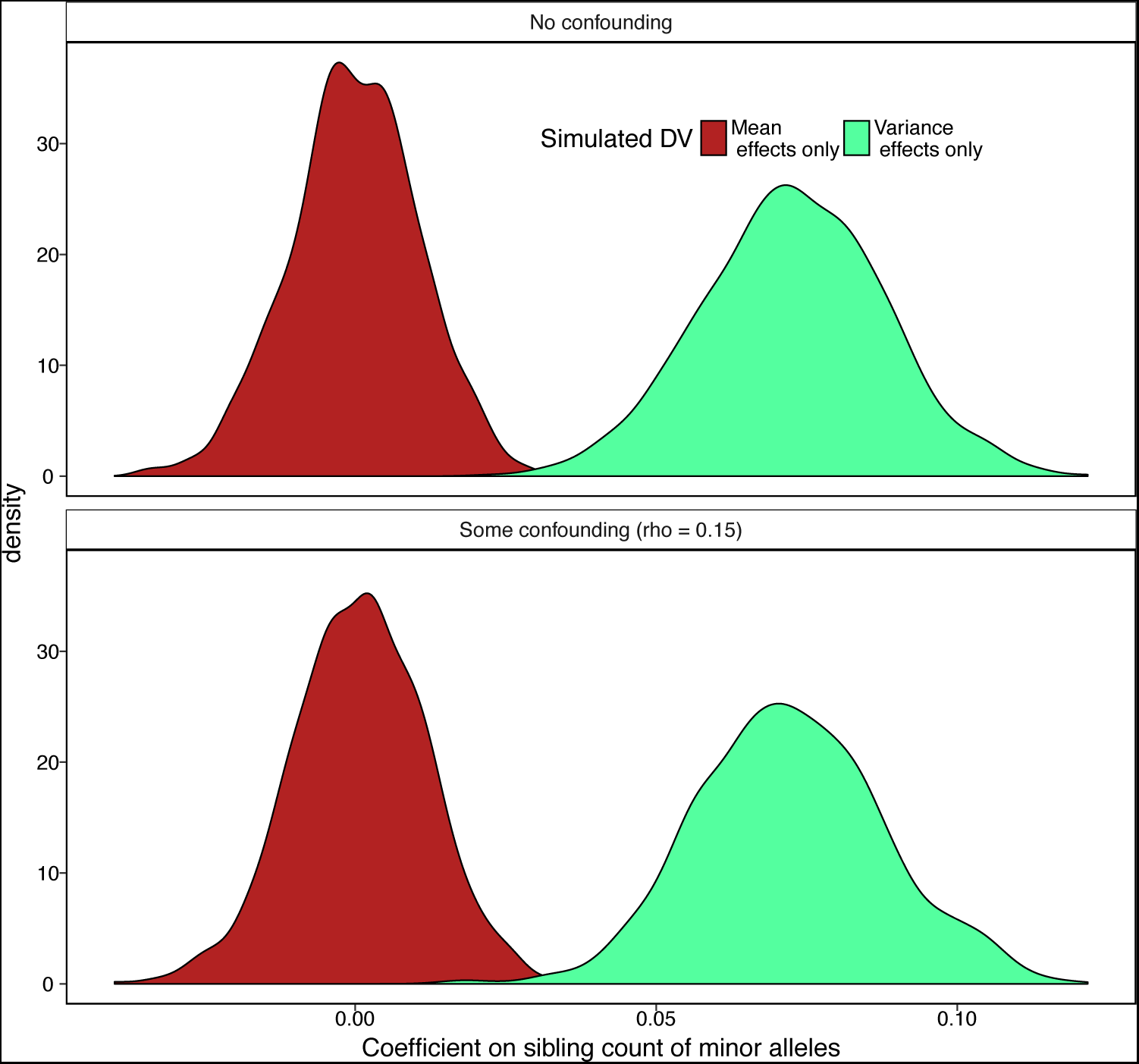
Results of sibling standard deviation method across 1000 replicates. The figure shows that both in the presence and absence of family-level confounding between the genotype and outcome variable, the method, which examines the effect of an additional minor allele in the sibling pair on the trait’s standard deviation, correctly estimates no variance effects (*β* = 0) when the outcome is simulated to have mean effects only, and correctly detects variance effects (*β* ≠ 0) when the outcome is simulated to have variance effects only.

S10 Table summarizes the percentage of simulations that reject the null of *β* on minor allele count equaling zero. In the presence of *no unobserved confounding*, the sibling SD approach has the expected type I error rate of correctly failing to detect 95.2% of variance effects when only mean effects are present (S10 Table). In contrast, the squared Z-score method correctly fails to detect 93.5% of variance effects when only mean effects are present. When we include a control for parental genotype in the *no unobserved confounding* case, this lower type I error rate comes at the cost of higher type II error, with the sibling SD method correctly detecting 92% of true variance effects across the replicates, versus closer to 100% for the other methods.

In the presence of *some unobserved confounding*, the sibling SD method greatly outperforms the other approaches in controlling type I error rates. While DGLM and the Squared Z-score method detect mean effects when only variance effects are present in 20-24% of simulations depending on whether ancestry controls or included and whether the data are demeaned, the sibling SD method detects mean effects when only variance effects are present at the expected rate of 4.8% (control for parental genotype) to 5.2% (no control for parental genotype) of cases. In addition, the method correctly detects variance effects when these effects are present in 91.6% (parent control) to 99.7% (no parent control) of simulations.

## Empirical application to height and BMI

### Sibling SD results: height and BMI

The simulation results show that in the absence of an unobserved confounder, the sibling SD method performs equally well as existing approaches to variance detection (squared Z-score; DGLM); in the presence of any unobserved confounder, the sibling SD method performs significantly better than these two approaches in correctly failing to detect variance effects when only mean effects are present. We now apply this method to two phenotypes: height and BMI. When we use data from quartets in the FHS and control for mean sibling-pair height, sex, mean pair age, within-pair age difference, and parental genotype in genome-wide regressions on the standard deviation of the sibling pair height we find four SNPs that are genome-wide suggestively significant (p<10-5) and meet other Hardy-Weinberg equilibrium (HWE) and minor-allele frequency (MAF) controls (see Methods): rs2804263 (MAF 30.8%); rs2073302 (MAF 39.1 %); rs8126205 (MAF 37.1 %) and rs4834078 (MAF 24.0 %) (Fig 3). For BMI, there were two SNPs that meet genome-wise suggestive significance: rs30731 (MAF 48.7%) and rs41508049 (MAF 10.3%)(Fig 4, with a regional linkage map of the SNP that replicates in S2 Fig). Our method, as expected, controls well for population structure: QQ plots for the height and BMI p-values do not show the telltale “early liftoff” typical of failure to control this confounder (S3 Fig).

**Figure 3.**
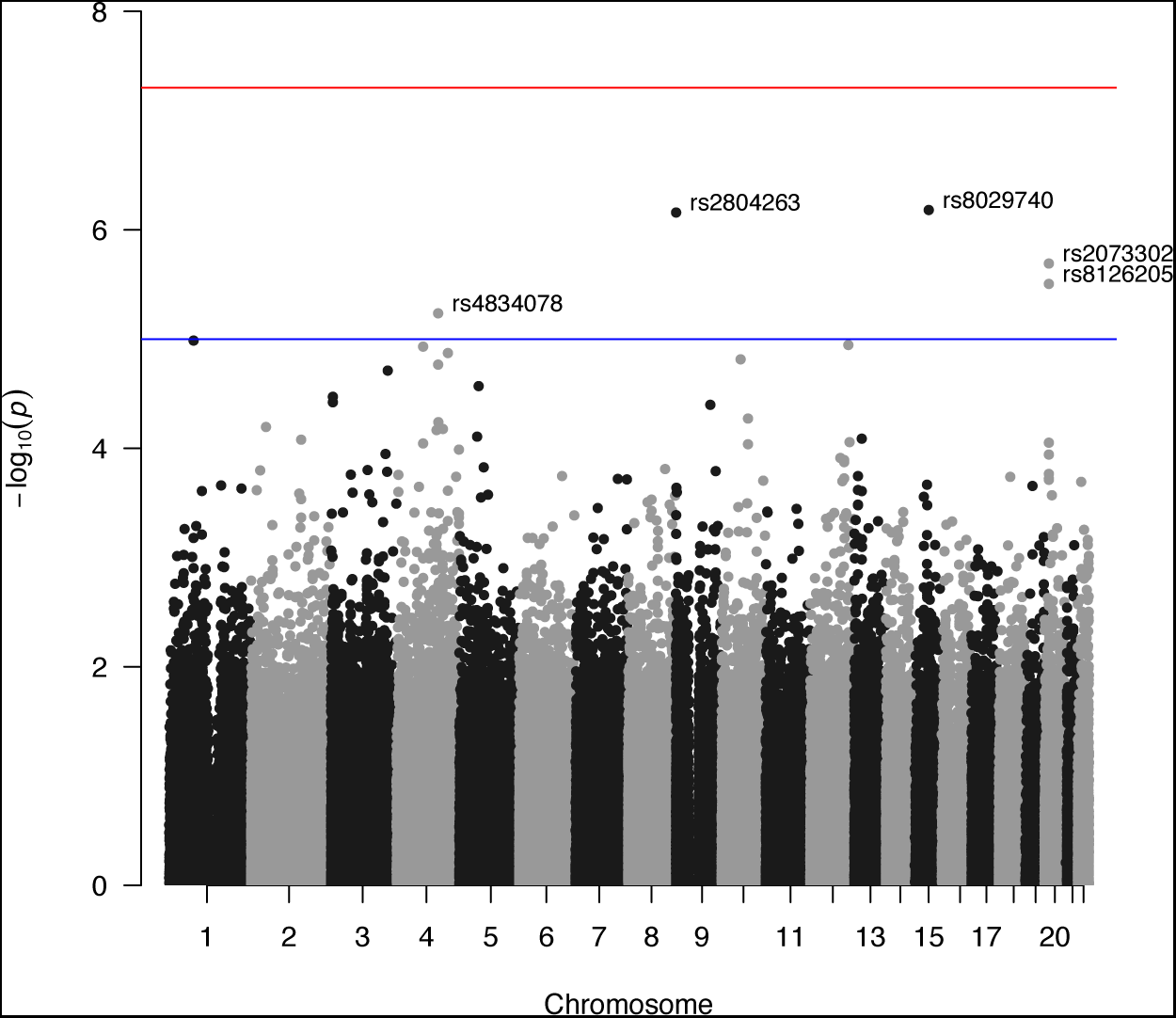
Manhattan plot of sibling variation in height among FHS 3rd generation sibling pairs. Results for the pairwise sibling standard deviation in height regressed against the sibling-pair minor count of alleles with controls for sex of sibship, mean age of siblings, age difference of siblings, sibling mean height, parental genotype.

**Figure 4.**
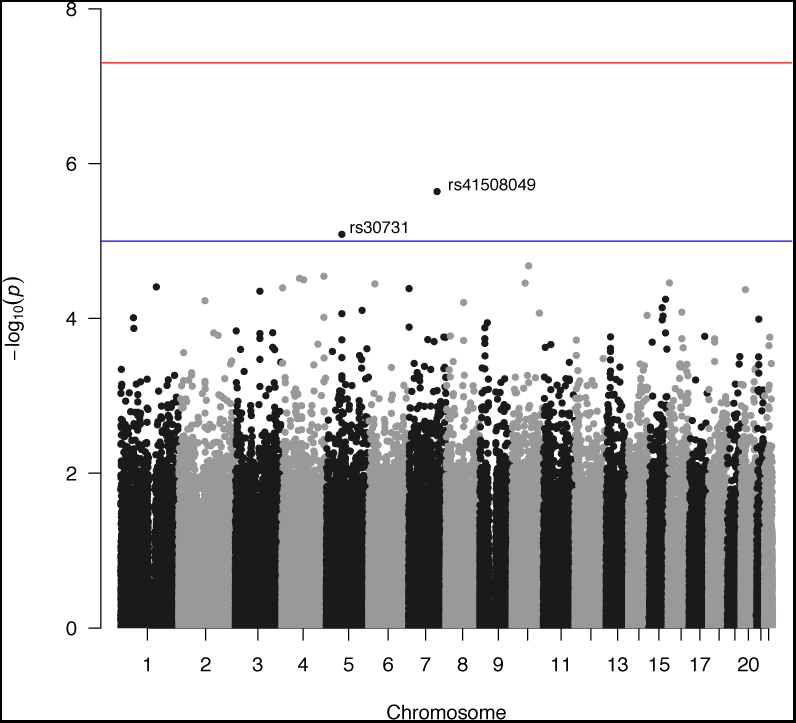
Manhattan plot of sibling variation in BMI among FHS 3rd generation sibling pairs. Results for the pairwise sibling standard deviation in BMI regressed against the sibling minor allele count with controls for sex of sibship, mean age of siblings, age difference of siblings, sibling mean BMI, and parental genotype.

The absence of genome-wide significant SNPs but presence of genome-wide suggestive SNPs may indicate that the method’s statistical power is low. A power analysis for the sample size utilized in the discovery dataset indeed supports this suggestion. For example, a single SNP would need to explain over 4.8% of the variation in the trait for the study to achieve 80% statistical power at the FHS sample size and a significance threshold of *p* < 10^−5^ (S4 Fig). An effect size of this magnitude is not expected for human complex traits (e.g., the largest effect of a single SNP for mean height explains approximately 1% of the variation). Power is very low near *R*^2^ = 0.01 (S4 Fig) in this particular sample. However, the figure shows how newly-released samples are adequately powered to detect the effects and highlight the method’s potential utility for better-powered studies.

Apparent effects on variance can be generated if effects of alleles at a locus are not additive (i.e., there is dominance) or if means and variances are correlated. Addressing the first issue, our approach does not control for non-additivity of allele effects at a locus, as it assumes a linear model. However, it does allow a test of whether an effect on variation net of mean was actually an artifact of non-linear effects on average rather than an actual variance effect. If the true relationship between phenotype and a sibling’s minor allele dosage were non-linear (i.e. revealed dominance effects) our initial findings could be entirely driven by divergence among those sibling pairs with two minor alleles. For example, if an individual with two minor alleles were significantly taller than an individual with either one or zero minor alleles (recessive effect) then when we collapsed the sibling pairs with two minor alleles, we could generate artifactual variation effects because among those sibships with two minor alleles, some would be distributed 0-2 (and thus one sibling would be taller than the other) while other sibships would be 1-1 (and thus would be the same height). Put together, it would appear that two minor alleles increased the variation net of mean effects. And if strong enough, such a misspecified effect could exert enough leverage to make a linear effect on variation appear across all allele numbers (zero to four for the sibship). These concerns appear not to apply to our analysis. A two-sample t-test of equality of means that compares homozygotes and heterozygotes for each significant SNP on the respective trait finds no differences in levels for either trait at the *p* < 0.05. Fig 5, which presents the mean and standard errors, shows the lack of significant differences.

**Figure 5.**
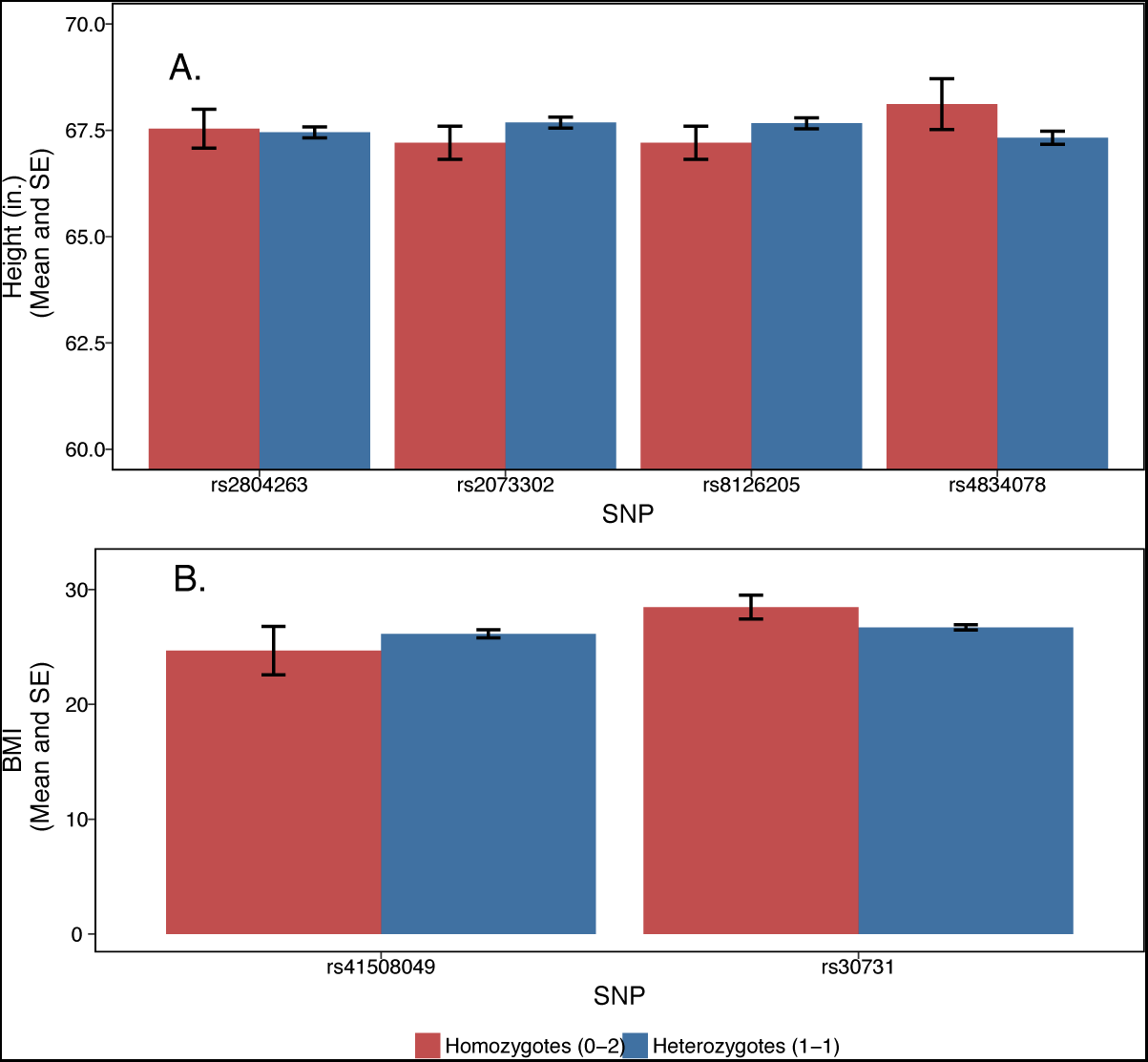
Test for spurious association with variance due to non-linear effects on mean levels. Mean and standard error for height (inches) and BMI among with two minor alleles is shown separately for homozygotes (one sibling with zero minor alleles and the other sibling with two) and heterozygotes (each sibling has one minor allele), for each genome-wide suggestively significant SNP for the respective trait (**A.** height; **B.** BMI). One significant SNP for height (rs8029740) is not depicted because there is only one sibling pair with the 1-1 allele combination and 0 sibling pairs with the 0-2 combination. A two-sample t-test for equality of means, estimated separately for each SNP, revealed no significant differences between the two groups for the top hits for each trait.

For the second issue, we can investigate correlated mean and variance effects in the present data. Although there is no overall correlation between effects on mean and on variance (as measured by squared Z-score and as previously reported ( [25]), the absence of the correlation appears to be caused by the fact that the vast majority of SNPs affect neither mean nor variance and therefore noise swamps out any signal; restricting the analysis to the SNPs that are the top hits reveal a very strong correlation between mean and variance effects for BMI (S5 Fig). To test these in an independent sample, the top SNP for variance in BMI found by Yang et al. ( [25]) shows clear association with mean BMI in the same data set from the GIANT consortium (S6 Fig), whereas the four suggestively significant SNPs in our analysis do not show an association with mean height in the GIANT consortium data (S7 Fig). Further highlighting the importance of our regression-based control for sibling-pair mean are the observations that: 1) sibling-pair standard deviation shows a strong positive correlation with sibling-pair mean, and 2) this correlation is not eliminated by using the sibling-pair coefficient of variation rather than the standard deviation (S8 Fig).

For replication analysis, we used respondents from the MTFS. (For list of proxy MTFS SNPs with information on MAF and linkage with FHS SNPs see S11 Table.) These families included phenotypic and genotypic information on pairs of twins as well as their parents, allowing us to replicate the sibling-based analysis with parental genetic controls so as to mimic random assignment of alleles. The MTFS has both dizygotic (DZ) and monozygotic (MZ) twins. Because MZ sibships do not vary in terms of cryptic genetic variation and may experience much more similar environments to each other than do genetically distinct siblings, we also repeated our replication analysis only with DZ twin sets but found that exclusion of MZ twins did not affect results. Another concern is that twins (even DZ twins) may experience more similar environments than singleton siblings; thus, our replication analysis may suffer from attenuation bias to the extent that the cause of variation is environmental and not cryptic genetic differences (which should, by contrast, be equivalent for singleton full siblings and DZ twins). Among the SNPs that were genome-wide suggestively significant for the height analysis, only two of the four had viable proxy SNPs in the MTFS dataset after quality control: rs2804263 and rs4834078 both had proxies whereas rs2073302 and rs8126205 did not (S11 Table). When we ran the analysis for the proxy SNPs in the MTFS dataset, none achieved statistical significance. Among the SNPs that were genome-wide suggestively significant for the BMI analysis, both SNPs had viable proxy SNPs in the MTFS dataset after quality control (S11 Table). When we ran the analysis for the proxy SNPs in the MTFS dataset, one SNP achieved statistical significance: rs30731 (proxy in MTFS: rs28636).

### Investigating rs30731/rs28636 for within-sibling variation in BMI

rs30731/rs28636, a SNP that significantly affects within-sibling variation in BMI and that replicated in the MTFS sample, is located on the MAST4 gene, which encodes a member of the microtubule-associated serine/threonine protein kinases. The proteins in this family contain a domain that gives the kinase the ability to determine its own scaffold to control the effects of their kinase activities.

GWAS studies have uncovered several significant associations between other SNPs on this gene and traits ranging from BMI to autism/PDD-NOS (S12 Table). One exception is a GWAS of childhood obesity conducted by the Early Growth Genetics Consortium ( [19]). The study found a genome-wide suggestively significant hit for rs28636 in the discovery sample that did not replicate. This non-replication could be due to the SNP being a variance-affecting locus that might show up in estimation of mean effects.

### Pathway and gene set analyses for all significant SNPs

In addition to investigating the gene function for the replicated SNP, we also performed two analyses that pool SNPs: gene-based and pathway-based pooling (see Methods). For the gene analysis, we found using PASCAL that the gene on which the significant SNP for BMI variability was located (MAST4) was significantly enriched (*p* = 0.0015 in the FHS data; *p* = 0.1 in the replication data). S13 Table shows other significant gene sets identified using PASCAL that replicated at various p-value thresholds in the MTFS data. VEGAS1 yielded no further significant gene sets.

i-GSEA4GWAS and PASCAL for pathway analysis each yielded some pathways that appeared to be significantly enriched (Table 2). One of these pathways, associated with within-sibship variance in height in the FHS data, replicated in the MTFS data: HSA04540 Gap Junction (*p* = 0.002 for each data set)(S9 Fig). HSA04540 includes members of several signaling pathways, including growth factors and their receptors, although any connection between these factors and organismal growth, as manifested in ultimate height, remains to be determined. Importantly, this pathway was not significant in the GWA for mean levels effects in either dataset. S14 Table shows replicated pathways for BMI and height variability estimated using PASCAL.

**Table 2.** Enriched canonical pathways for height and BMI sibling-pair standard deviations in FHS, estimated using i-GSEA4GWAS

| Pathway/Gene set name | P-value | FDR | Sign. Genes/<br>Slctd. Genes/<br>All Genes |
| --- | --- | --- | --- |
| **HEIGHT** |  |  |  |
| GLYCEROLIPID METABOLISM | < 0.001 | 6.0000000000000001E-3 | 20/35/45 |
| HSA00561 GLYCEROLIPID METABOLISM | < 0.001 | 8.9999999999999993E-3 | 27/49/58 |
| VEGFPATHWAY | < 0.001 | 1.5666665999999999E-2 | 47046 |
| HSA03030 DNA POLYMERASE | 1E-3 | 2.76E-2 | 45920 |
| BILE ACID BIOSYNTHESIS | 3.0000000000000001E-3 | 3.175E-2 | 46712 |
| HSA04540 GAP JUNCTION* | 2E-3 | 4.1666667999999997E-2 | 42/78/98 |
| HSA00564 GLYCEROPHOSPHOLIPID METABOLISM | 5.0000000000000001E-3 | 0.24111110999999999 | 20/49/68 |
| **BMI** |  |  |  |
| HSA00591 LINOLEIC ACID METABOLISM | < 0.001 | 6.0000000000000001E-3 | 11652 |

\*Replicates in MTF5 data

## Discussion

Our analysis extends earlier work that aimed to map variance-controlling loci in humans ( [25]). Although the prior work enjoyed greater statistical power, it also had more potential for bias-due both to environmental confounding and to conflation of mean and variance effects. Indeed, Yang et al. identified a locus regulating BMI variability that is also strongly associated with mean levels and for which a gene-by-environment interaction effect on mean has been shown. In the present study, we first show via simulation that two widely-used methods for detecting variance-effecting loci-the squared Z-score method and DGLM-fail to adequately distinguish between a locus affecting the trait mean and a locus affecting the trait variance in the presence of unobserved confounding. The sibling SD method, by controlling for the sibling mean of a trait and identifying off of random, between-sibling variation in allele counts, does distinguish between these effects. Applying the method to data, we were able to perform within-family analysis on two samples of white Americans, completely free of population stratification, largely devoid of rGE confounding, and with controls for mean level effects as well as checks for non-linear (i.e. dominance) effects on mean levels. One SNP for BMI variability, located on the MAST4 gene, replicated. Notably, the SNP, unlike many on FTO, has not been found to affect mean levels of BMI. Like the latest methods to map variance-controlling loci in controlled crosses ( [64]), our approach therefore avoids common confounds. At the same time it overcomes problems specifically associated with human traits, including the construction of variance-affecting environments, that existing regression-based methods for detecting vQTLs fail to address because they allow controls solely for observed confounders (e.g., [11,12,24,64]).

Though underpowered in the FHS and MTFS sample sizes used in the present analysis (S4 Fig), our results strongly support the benefits of approximating a randomized genetic experiment by analyzing within-family variation while controlling for parental genotype. Such an analysis addresses the possibility that it is merely cross-family environments interacting with a mean effect and/or population structure that produce apparent association with variability. Meanwhile, parameterizing the estimand as spread (SD) net of sibship mean levels provides a robust, flexible way to conceive of variation—that is, rather than parameterizing the relationship between mean and variance a priori by using the coefficient of variation or some similar summary statistic. The trade-off inherent to our approach is that environmental and phenotypic variation within sibships may be attenuated, reducing statistical leverage; the extent of such a dynamic is wholly dependent on phenotype, of course. Although the within and between family components of the variation in the phenotype can be measured to determine whether or not the phenotype is suitable for such an approach, the extent of variation within and between sibships in the unmeasured environmental factors that matter is, of course, unknown.

Moving forward, we see three applications of the method: (1) combining the method with twin studies to better distinguish between *GxG* versus *GxE* effects that each contribute to trait variability; (2) examining the heritability of plasticity in a trait as a supplement to examining the heritability of levels of a trait; and (3) generating weights for polygenic scores to predict trait variance. We discuss each in turn.

### Variability across MZ versus DZ twins

First, we can combine the present within-family approach to measuring phenotypic variability with the classic twin comparison approach of behavior genetics, we obtain a method to distinguish between GxE and GxG interaction effects that may be revealed in a vGWAS for variation regulating loci. Namely, if a particular allele produces more variability among dizygotic twins than among monozygotic ones, we can infer that the difference between those allelic effects is attributable to two forces: 1. Putatively greater environmental differences within DZ twin pairs than within MZ twin-ships; and/or 2. The greater (cryptic) genetic variation within DZ pairs as compared to their MZ counterparts. Since prior work ( [18]) shows that the equal environments assumption seems to hold for a wide range of outcomes, thus weakening support for 1 as the explanation, we can attribute the bulk of the difference in trait variation between DZ versus MZ twins to the theory that the allele in question is not only buffering the environment but also serving as a phenotypic capacitor (i.e. repressing cryptic genetic variation).

### Estimating the heritability of plasticity

Another way to combine the present approach with classical statistical genetic techniques is to supplement estimates of the heritability of *levels* of a trait with estimates of the heritability of *plasticity* in a trait. For instance, GREML involves partitioning the observed phenotypic distance in a trait between individuals into the sum of genetic and environmental contributors to this distance. If we switch from individuals as the unit of analysis when measuring this distance to sibling pairs as the unit of analysis, we can calculate the phenotypic distance between siblings in each pair and the average genotype at each locus across the two siblings. The, we can place unrelated sibling pairs in a Genetic Relatedness Matrix and contrast the genetic distance between sibling pairs to the amount of variability for the phenotype the pairs display to recover estimates of the heritability of this variability.

### Constructing vPGS

Currently, researchers develop and use polygenic scores (PGS) that predict mean levels of a trait. We can extend the polygenic score approach to develop scores that predict variance in a trait (vPGS). Coefficients for a vPGS construction in a prediction sample would be obtained from a vGWAS done within families with sibling sets in the discovery sample to obtain estimates that better distinguish between mean and variance effects, but the application of coefficients could then be to the individual person. S4 Fig shows that the method is adequately powered to detect effects of SNPs that explain fewer than 1% of the variation in traits in samples like the UK Biobank that could be used at this discovery stage.

Having a polygenic risk score that predicts particular forms of phenotypic variability may be helpful for researchers hoping for non-null results with respect to a given phenotypic measure who are therefore looking to recruit sensitive subjects for experimentation that involves specific environmental exposures. Per the earlier discussion, if calculated from pairs of MZ twins, such a polygenic score would capture only environmental sensitivity. But if a vGWAS of MZ twins and DZ twins were conducted, the results could be differenced out to provide a measure of phenotypic capacitance-i.e. regulation of internal, genetic variation.

Beyond telling experimenters which subjects may be more genetically sensitive, such a phenotypic capacitance score may have important predictive power in terms of disease. Namely, cancer, autoimmune diseases, metabolic syndrome and other irregularities of cell or system stability may themselves be predicted by a genetic architecture that is less robust. Thus, a genetic screening for a tendency toward developmental plasticity (i.e. if the plasticity score was calculated on developmental indicators such as height) may be diagnostic. If applied to behavioral phenotypes, such a score could be predictive of mental disorders that reflect a lack of canalization of mind, so to speak, such as schizophrenia.

In light of this discussion, we think that there is benefit to combining prior, pedigree-based approaches with newer GWAS methods to better estimate variance effects (as well as levels effects). Thus, we recommend that consortia of cohorts with genome-wide data on sibling pairs at the minimum, quartets ideally, be formed to advance GWA to a more solid foundation of inference that approximates the unbiased estimates of lab-based genetic manipulations by taking advantage of random differences in sibling genotypes.

## Materials and methods

### Simulation study

#### Generating genotype and trait data

The simulation proceeds in four steps. First, we generate genotypes for parents and offspring. Second, we generate an unobserved family-level confounder that is correlated to varying degrees with the observed sibling genotype. Third, we use the genotype to generate four traits:

1. Trait with neither mean nor variance effects
2. Trait with mean effects but not variance effects
3. Trait with variance effects but not mean effects
4. Trait with both mean and variance effects

In this third step, we generate three versions of each of the four traits: 1) a version that is not affected by between-family confounding; and a version that is affected by 2) moderate levels of between-family confounding and 3) high levels of between-family confounding. Fourth, we explore how the process in steps one through three generates between-family confounding that is not fully addressed by controls for the subpopulation that generates the genotype. Finally, we compare the performance of the sibling standard deviation approach to other methods. We describe the first four steps in the present section, and summarize the results of step five in the results.

All steps were repeated for 1000 replicates of size 8000 (4000 sibling pairs).

#### Step one: generating genotypes

We use the following process to generate genotypes for parents and offspring, repeated separately for each of the 1000 replicates:

1. *Generate parent genotypes:* we use the function `simMD` within R’s `popgen` package to generate parent genotypes, and use the following parameters in the present simulation:
  - *N* = 8000 parents, *N* = 4000 families with 2 offspring per family
  - 4 subpopulations, with *c* = 0.01 representing the extent to which each subpopulation differs in allele frequencies of SNPs from typical values
  - 1 causal snp. Traits with no effects were thus generated by setting the coefficient on the parameter that governs the relationship between 1) the allele on trait mean, and 2) the allele on trait variance, to zero.
  - Allele frequency (*p*) of each SNP randomly drawn from a uniform distribution with bounds at [0.1,0.9]
2. *Generate offspring genotypes* : we use the parent genotypes generated in step 1 to generate offspring genotypes assuming random mating and segregation
3. Step one and step two result in parent and offspring genotypes we use in steps two and three of the simulation

#### Step two: generating a family-level confounder correlated with genotype

Consider the following model for the relationship between a SNP and phenotype for an individual *i*, nested in family *j*, for snp *k*. For now, we just consider identifying the causal effect of an allele on the mean of *Y* :

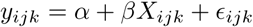

Linear regressions such as those run in GWAS, which often restrict the sample to unrelated individuals, ignore the grouping structure of the family and estimate the following model:

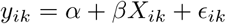

Random effects models acknowledge this grouping structure by positing that each family has its own intercept that shifts the outcome up or down. The random effects models then estimate these *α_j_* using a distribution that pulls some of the family-specific intercepts 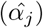 towards the mean intercept across families (*μ_α_*) depending on *σ_α_* ( [31]):

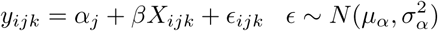

Random effects methods generate unbiased estimates for *β* (the causal effect of an allele on a trait) when there is no unobserved confounding at the family level that is correlated with genotype of other observed covariates (*cor*(*X_ijk_*, *α_j_*) = 0))( [21]). However, these methods generate biased estimates when *cor*(*X_ijk_*, *α_j_*) ≠ 0). To generate simulations to test this bias in the genetics context, we use the following process to generate a family intercept that is correlated with offspring genotype, and through the process described in step three, is also correlated with the trait:

1. Operationalize “genotype” as the sibling pairs’ summed minor allele count at that locus
2. Choose a *ρ* parameter for the degree of correlation, and construct a variance-covariance matrix *Q* representing the correlation between the genotype and family-level intercept. E.g., if *ρ* = 0.3:

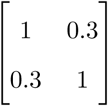
3. Take the cholesky decomposition of *Q*
4. Generate a starting value for the family-level intercept that is not correlated with genotype (*α_j_* ~ *N*(0,1)); this starting value provides a baseline set of intercepts uncorrelated with genotype that we then rescale to be correlated with genotype to varying degrees.
5. Multiply cholesky decomposition by matrix containing the genotype and the non-correlated intercept to generate *α_j_* correlated with the sibling pair’s genotype
6. Repeat for three values of *ρ*:
  1. *ρ* =0
  2. *ρ* =0.15
  3. *ρ* =0.3

#### Step three: generating traits

To generate traits, we used a similar process for simulating traits as used in [11]. We generated four general types of traits (traits with neither mean nor variance effects; traits with mean effects only; traits with variance effects only; traits with both mean and variance effects) using the following general setup, and varying the *γ* and *ɛ* parameters, where *i* represents an individual and *k* indexes a SNP. Sex was simulated from a binomial distribution with *p* = 0.5. Age was simulated from a normal distribution ~ *N*(*μ* = 50, *sd* = 10). *G*_1_ indicates heterozygotes, while *G*_2_ indicates minor allele homozygotes. Across all simulations with mean or variance effects, an additional minor allele results in *increases* in the mean or in the variance:

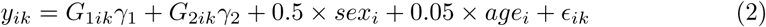

1. Neither mean nor variance effects:
  - *γ*_1_ = *γ*_2_ = 0
  - *ɛ* ~ *N* (0,1)
2. Mean effects only
  - *γ*_1_ = 0.15; *γ*_2_ = 0.35
  - *ɛ* ~ *N*(0, 1)
3. Variance effects only
  - *γ*_1_ = *γ*_2_ = 0
  - *ɛ* ~ *N*(0,1) for major allele homozygotes; *ɛ* ~ *N*(0,1.15^2^) for heterozygotes; *ɛ* ~ *N*(0,1.4^2^) for minor allele homozygotes
4. Mean and variance effects
  - *γ*_1_ = 0.15; *γ*_2_ = 0.35
  - *ɛ* ~ *N*(0,1) for major allele homozygotes; *ɛ* ~ *N*(0,1.15^2^) for heterozygotes; *ɛ* ~ *N*(0,1.4^2^) for minor allele homozygotes

For the simulation to address the possibility of confounding by unobserved, between-family factors, we also modify equation 2 to include the family-level intercept that is correlated with observed genotype at varying levels (no correlation, medium correlation, high correlation), and refer to the latter two outcomes as “confounded outcomes”, where *i* refers to an individual and *j* indexes the family that the sibling pair was generated from:

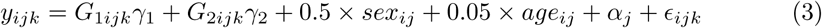

#### Step four: controlling for ancestry

To control for ancestral background in the methods that follow, we control for an indicator for which of the four subpopulations generated the individual/pair’s genotype. We used an indicator for ancestry rather than ancestry derived from genotype because, and following the general simulation method used in [11] where traits were generated using one SNP because each SNP-trait association is estimated separately, the genotype has only SNP.

As the strength of between-family confounding increases, the correlation between these indicators for population stratification and the family-level intercept increases. To illustrate this increase, we run the following regression for each of the three degrees of correlation between family-level intercepts and observed genotypes:

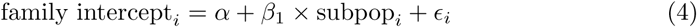

We then calculated the percent of *β*_1_ ≠0 with *p* < 0.05 across the 1000 replicates with that simulated level of correlation. S15 Table summarizes the results and shows that as family-level confounding increases, the variable for population stratification better predicts this confounding. However, as we show in the results that follow, while this correlation between the population stratification indicators and family-level confounding *reduces* bias in estimates that this confounding causes, the control does not fully eliminate bias.

Equation 4 looks at the strength of relationship between ancestral background an the family intercept at various degrees of confounding, and shows how this strength increases as the degree of confounding increases. The reason this confounding biases estimates of the effect of the minor allele count on a trait is because this family intercept is also correlated with genotype. To show this, we run the following regressions at each level of confounding:

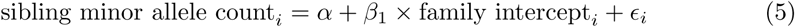

S16 Table presents the mean *β*_1_ across each level of confounding, and shows that as confounding increases, the relationship between family intercept and the minor allele count increases. In the presence of zero between-family confounding, the slightly positive *β*_1_ is caused by the correlation between parent and offspring genotypes that generates correlated genotypes between siblings of the same parents.

### Power

For the trait simulated to have variance effects, the average effect size was *R*^2^ = 0.01. With the *N* = 4000 sibling pairs sample sized used in the analysis, the power to detect these effects approaches 100% (99.9%). As a result, the simulation results consistently detect variance effects when these are present and the comparisons mainly focus on when the different approaches-squared Z-score; DGLM; sibling SD-correctly fail to find variance effects when these are *not* present (type I error).

### Analysis

All analysis was performed using R. The following packages were used to estimate the models (see Results):

- Power analysis: `pwr` packages with `pwr.f2`.test command
- Pooled regressions: `lm` with default settings
- Random effects model: `plm` with model = “random”
- Fixed effects model: `plm` with effects = “within” and index of the family identifier
- DGLM: `dglm` with a gaussian-family link function and REML as estimation method
- Squared Z score method: after estimating the Z-score separately by sex, `lm` with default settings
- Sibling SD method: after estimating the sibling standard deviation, `lm` with default settings

### Empirical application

#### Data

Data for discovery analysis come from the Framingham Heart Study (FHS), second (parental) and third (sibling) generation respondents. (This dataset is publicly available through dbGaP http://www.ncbi.nlm.nih.gov/gap. QC code can be obtained from the FHS investigators ( [1]) The FHS is, in fact, one of the cohorts included in the GIANT meta-analysis performed by Yang et al. ( [25]). Height and weight were taken from clinical measurements and then BMI was calculated as (weight in kilograms)2 / height in meters. Genotypes were assayed using the Affymetrix GeneChip Human Mapping 500K Array and the 50K Human Gene Focused Panel. Genotypes were determined using the BRLMM algorithm. Our analysis began with the original 500,568 SNPs, and resulted in 260,469 SNPs available for analysis after cleaning (e.g., HWE screens and a MAF cut-off of 0.05). The screens were conducted using all available individuals with genetic data, not only those that were included in this analysis. Genome-region association plots were produced using SNAP ( [47]), except for those of published GIANT consortium data, which were produced using LocusZoom ( [60]). Regional linkage maps were produced using SNAP ( [47]) and data from the 1,000 Genomes CEU Panel ( [2]), which also provided the reference MAFs for S1 Table.

Among third-generation respondents, the numbers in our sample by sibship size are presented in Table 3. The 200 families with only one sibling in the data drop from the sibling analysis. Those with more than two contribute multiple pairs to the data; however, our final analysis selects only one pair per second-generation family as the more complicated error structure with multiple pairs leads to early takeoff on QQ plots. The siblings are genetically related but are not DZ or MZ twins.

**Table 3.** Distribution of third generation siblings included in data by sibship size:

| # 3rd-G Sibs in Family: | 1 | 2 | 3 | 4 | 5 | 6 | 7 | 8 | 9 |
| --- | --- | --- | --- | --- | --- | --- | --- | --- | --- |
| N (pairs): | – | 292 | 483 | 504 | 250 | 150 | 189 | 28 | 36 |
| Families: | 200 | 292 | 161 | 84 | 25 | 10 | 9 | 1 | 1 |

Total actual N of sibling pairs after random selection of one per family: 583

The Minnesota Twin Family Study (MTFS) replication data were genotyped on the Illumina 660W Quad array ( [55]) and phenotypes can be found elsewhere ( [42]) Quality control procedures were applied separately to each individual cohort. Individuals with a call rate < 0.95 (*N* = 22), estimated inbreeding coefficient > 0.15 (N=2), or showing evidence of non-European descent from multidimensional scaling (N=298, mainly individuals with Mexican ancestry) were removed. Individuals were considered outlying from European descent if one or more of the first four eigenvectors were more than three standard deviations removed from the mean. SNPs with MAF < 0.01, call rate < 0.95 or HWE-test p-value < 0.001 were removed. The reason for the lower MAF threshold in the MTFS sample rather than FHS sample is that for the latter data, internal QC procedures remove SNPs at this threshold prior to releasing the data to researchers. For our analysis, we included both the MZ and DZ twin pairs because restricting to DZ twin pairs that more closely approximate the sibships in the FHS discovery sample does not change the substantive findings. Because the discovery sample and the replication sample were genotyped on different arrays, we deployed SNAP to find corresponding SNPs ( [47]). The resulting sample size for our analysis was 1,048 pairs.

In addition to the GWA analysis in the discovery and replication sample, we performed two sets of analyses that pooled SNPs across multiple software implementations. First, we investigated gene-based pooling using VEGAS1 ( [53]) with the following parameters: CEU subpopulation specified, no assumed allele frequency difference by sex, and using all SNPs (not just best hits) within the gene region itself (+/−0 KB). We estimated pathway-based pooling using i-GSEA4GWAS ( [85]). Because estimation of significant pathways and gene sets can be sensitive to different algorithms’ methods of computing p-values and controlling type I error rates, we test the robustness of these findings with PASCAL ( [50]). These analyses were performed for both BMI and height in both the discovery and replication sample. For the discovery sample, gene-based and pathway-based analyses were performed using 260,434 variants input; 239,526 variants used; 14,783 genes mapped; 221 gene sets selected. For the replication sample, gene-based and pathway-based analyses were performed using 522,726 variants input; 487,692 variants used; 16,840 genes mapped; 259 gene sets selected.

#### Statistical Analysis

All analysis was performed using R. The power analysis depicted in S4 Fig to estimate the power of the approach at varying putative effect sizes was performed using the *pwr* package in R [15], varying *n* to reflect the present sample sizes and recently-released samples with larger sibling populations and showing power separately for two p value thresholds (*p* < 10^−5^ for discovery analyses; *p* < 0.05 for replication). More precisely, we used the following procedure:

1. Vary the putative *R*^2^ explained by a SNP from 0.0 to 0.01, incrementing by 0.0001
2. Translate *R*^2^ into effect size 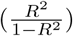
3. Since we are analyzing power to estimate the *β* in a linear regression, using the `pwr.f2.test` command to estimate power at varying effect sizes

Heteroscedasticity-robust standard errors should not substantially affect power under the present sample size ( [43]).

For the main analysis, sibling-pair standard deviations (SD) were fit by linear regression using the `lm` command with default options to the following model, where the key regressor is the number of minor alleles for the pair of siblings at a given locus. Because this number is for two individuals, the range is 0 to 4:

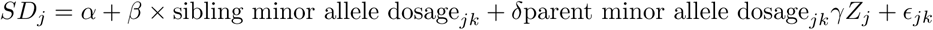

*j* indexes a sibling pair, *k* indexes a SNP, minor allele dosage is the total number of minor alleles in a sibling pair, parent minor allele dosage is the total number of minor alleles across the parents, *Z_j_* is a vector of sibling pair-level controls that includes controls for the mean level of the trait in the sibling pair, pair sex (MM or FM or FF), mean pair age, and the within-pair age difference, and *ɛ_jk_* is the residual for sibling pair *j* at snp *k*. As we show in the simulation study, the method returns similar results if parental genotype is or is not included. Qualitative results do not change if we instead specify the mother’s and father’s genotypes separately. In the model, *SD_j_* is the standard deviation of a trait within a sibling pair, calculated as follows, where *i* indexes an individual sibling in the pair, *j* indexes the pair, *x* refers to the trait, and 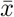 refers to the mean of the trait across the two siblings:

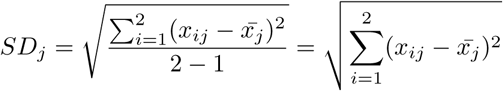

Huber-White standard errors robust to clustering on pedigree ID (to account for correlated errors among cousins: sibling pairs that share the same grandparents but not the same parents) were calculated for the FHS analysis in the following way, where *i* = sibling pair 1, *j* = sibling pair 2, *n* = total sibling pairs, *g* = pedigree grouping, and *k* = SNP. *σ* refers to the residual variance across the sample, and *σ_i_* refers to the residual variance for an individual.

For the simple case where 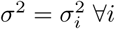 and 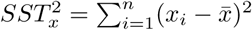:

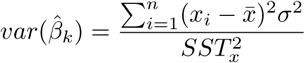

To account for errors that may be heteroskedastic and correlated within a shared pedigree, we adjust the variance to be robust to cases where 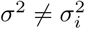 and where for *g* = *g′* (two sibling pairs share same grandparent/pedigree ID). 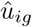 represents the observed residual for participant *i* belonging to pedigree *g*. For the case of pedigree ID’s with two or more sibling pairs, this becomes the following variance robust to heteroskedastic and correlated errors, with an indicator function for when the sibling pairs belong to the same pedigree:

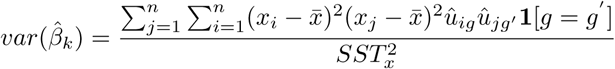

For the case of pedigree ID’s with one sibling pair only, the above equation reduces to the following variance robust to heteroskedastic errors ( [10]):

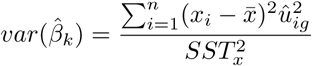

Which we can simplify further to:

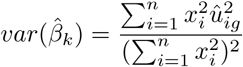

## Acknowledgements

For helpful feedback on this work, we would like to thank members of the Siegal Lab at the NYU Center for Genomics and Systems Biology as well as Justin Blau, Richard Bonneau, and Michael Purugganan of the NYU Department of Biology, Tomas Kirchoff of NYU Medical School, and Boriana Pratt of Princeton’s Office of Population Research. The authors are also grateful to participants in the University of Colorado’s 2012 workshop on Integrating Genetics and the Social Sciences for helpful comments.

## Supporting information

**S1 Fig.**
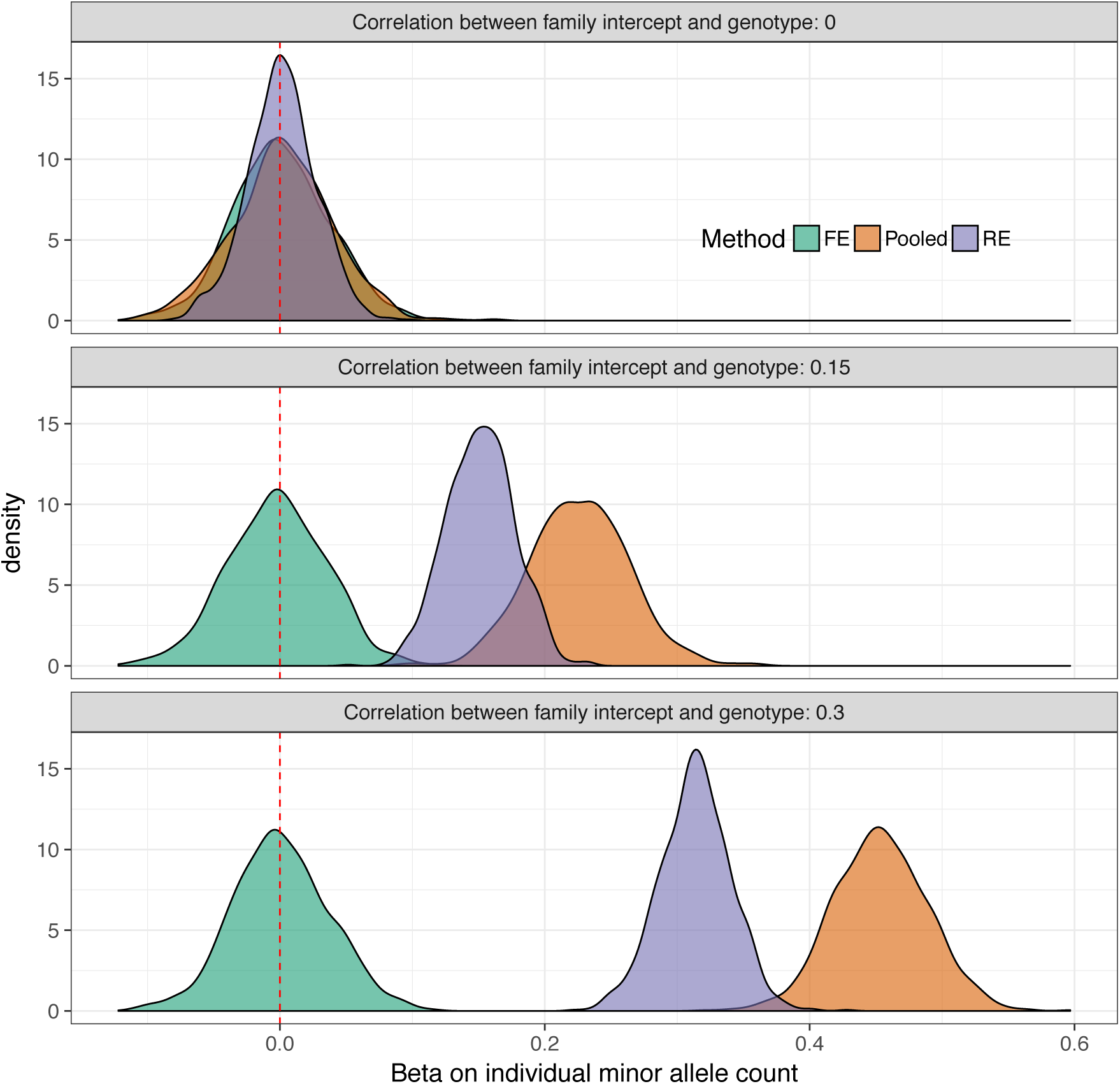
Estimated coefficients on SNPs for simulated dependent variable with *no effects* and confounding between a family-level indicator, genotype, and outcome. The red dashed line represents the true snp level effect (*β* = 0), while the density curves show the range of estimated 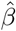 for each of the models. We see the fixed effects model correctly centers the 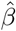 near the *β* = 0, while the other family-level random effects (random intercept) and pooled regression show estimates with significant upward bias in the presence of confounding. However, the random effects has the advantage of smaller sampling variance (more efficient estimator) across all levels of confounding because it pools estimates across families.

**S2 Fig.**
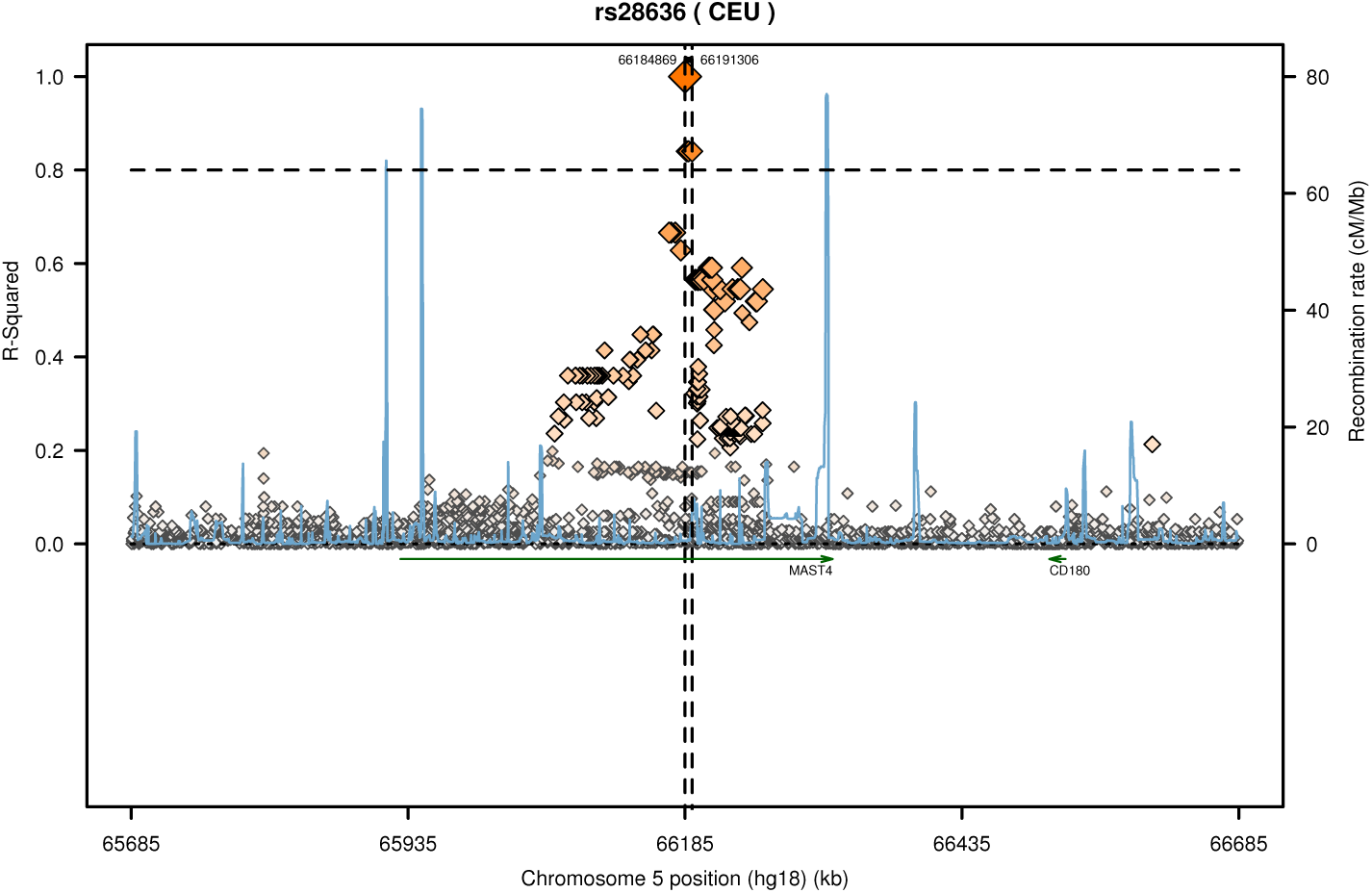
Regional linkage map for FHS genome-wide suggestive SNPs for sibling-pair standard deviation in BMI from 1,000 Genomes, CEU Panel. Maps produced by SNAP ( [47]).

**S3 Fig.**
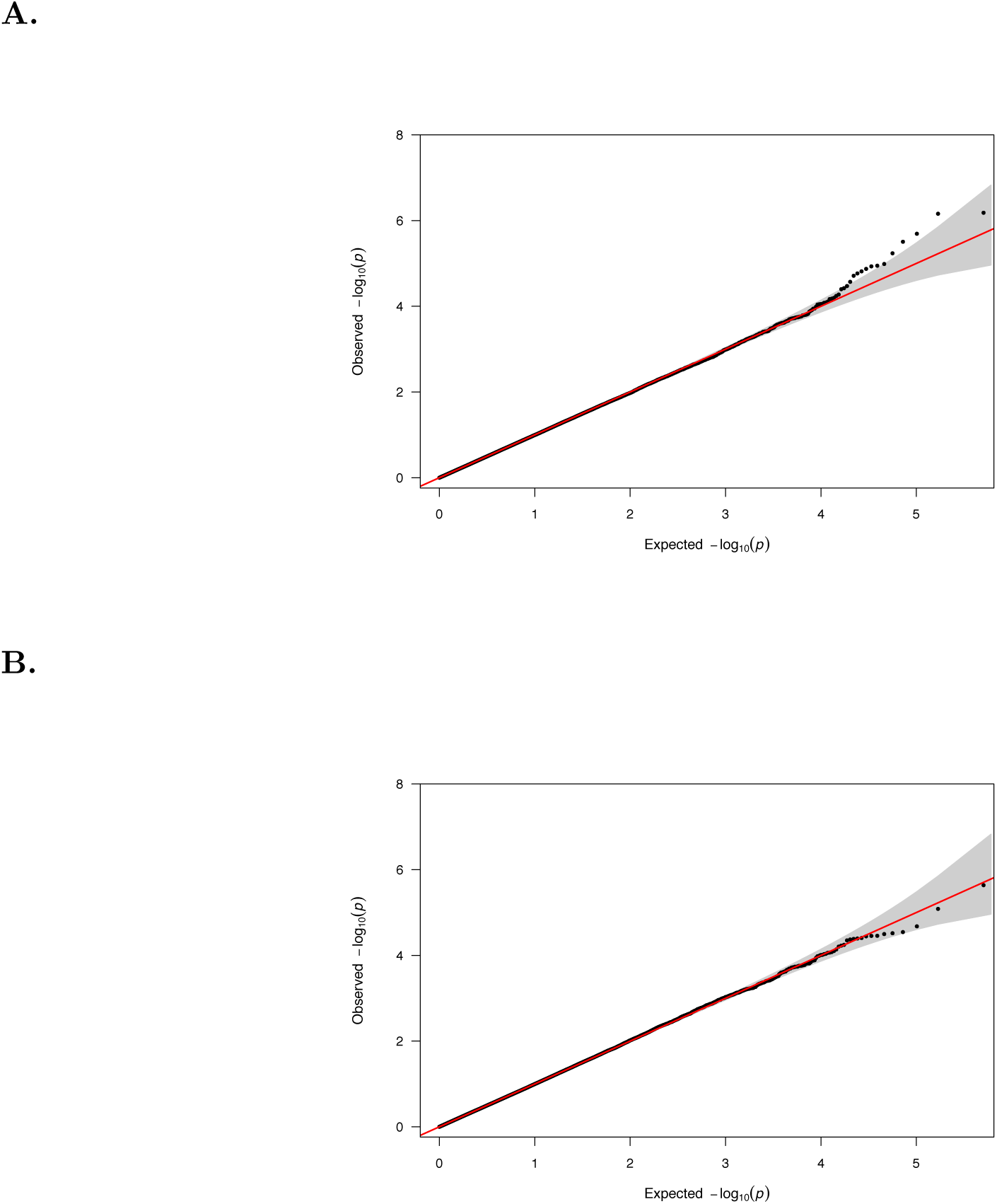
QQ plots associated with Manhattan plots in Fig. 1. A) Observed versus expected p-value distributions for analysis of sibling-pair standard deviation in height for FHS generation-three respondents with controls for parental genotype, mean height of sibling pair, sex, and sex difference. B) Same as in (A) except for BMI instead of height. Shaded gray regions depict 95% confidence intervals.

**S4 Fig.**
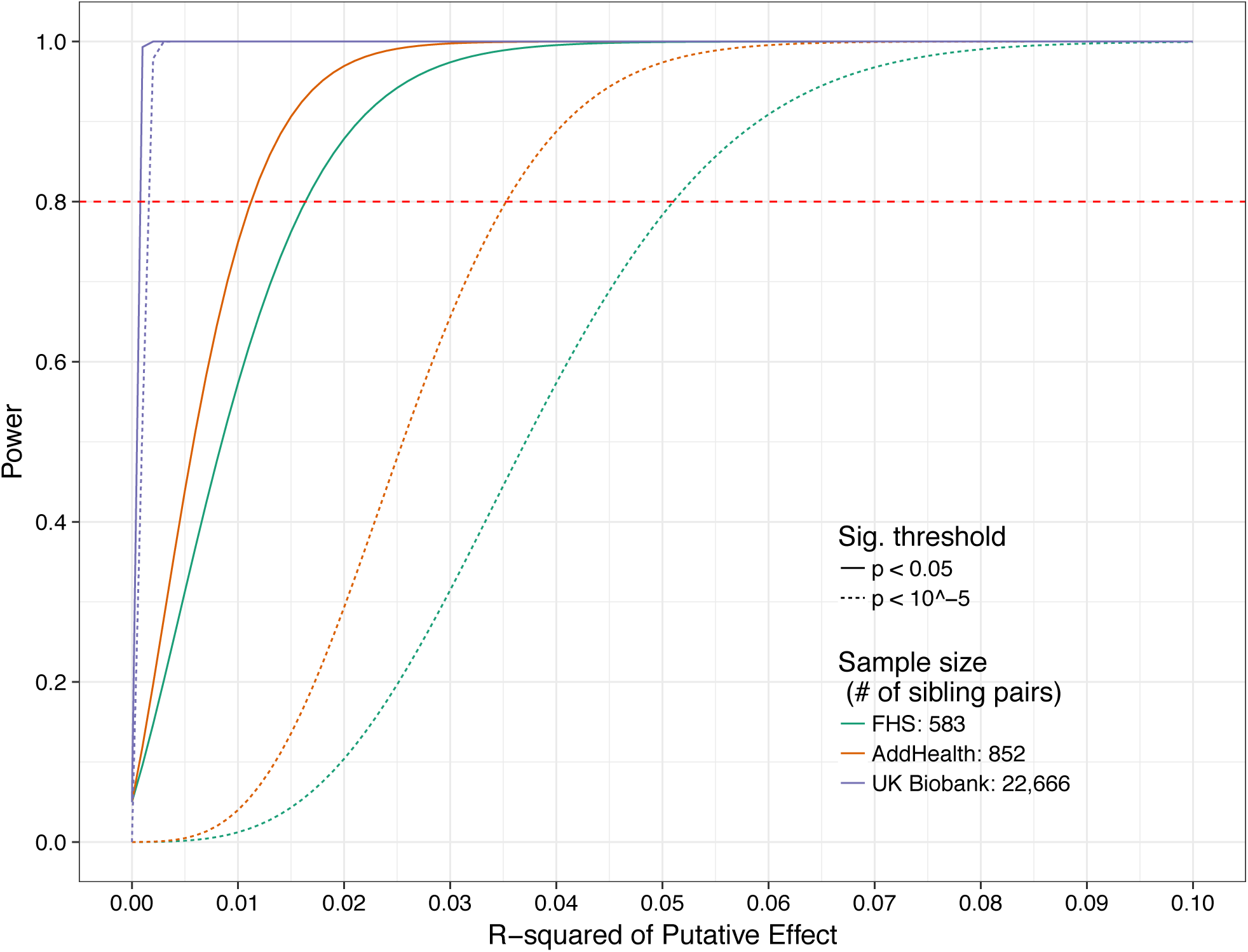
Power to detect an effect size of *R*^2^. The figure contrasts power at three potential sample sizes (defined as the number of sibling pairs in the data) (see Methods): 1) the Framingham Heart Study (FHS) sample used in the present analysis; 2) the Adolescent and Longitudinal Study of Health (AddHealth) sample; and 3) the UK Biobank sample. Likewise, the figure contrasts two potential p-value thresholds: *p* < 10^−5^ for the discovery analysis; *p* < 0.05 for the confirmation analysis. The figure shows that although the sample used in the present analysis (FHS) is not adequately powered to detect realistic effect sizes of *R*^2^ < 0. 01, newly-released datasets with larger sibling subsamples are adequately powered to detect effects using the method.

**S5 Fig.**
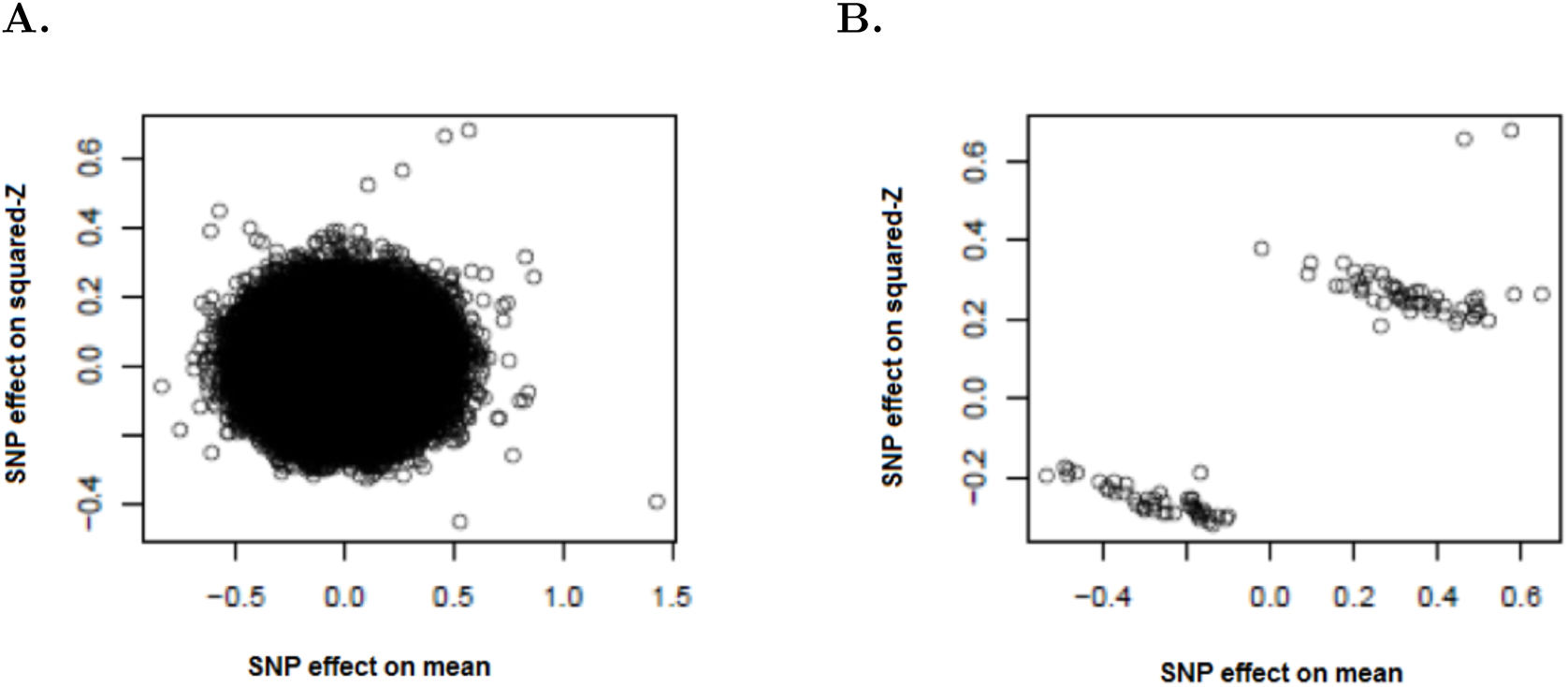
Correlation between SNP mean effects and SNP association with squared Z-scores. SNPs are normalized for minor allele frequency (W). A) For each SNP, association between the SNP and squared Z-scores for BMI is plotted against the SNPs effect on mean BMI (correlation approximately zero). B) Same as in (A) except only the top 100 SNPs (based on mean effects on BMI) are shown (correlation 0.87). This shows that SNPs that have significant mean effects on BMI have effects on the variance that are significantly correlated with the SNP’s effects on the mean.

**S6 Fig.**
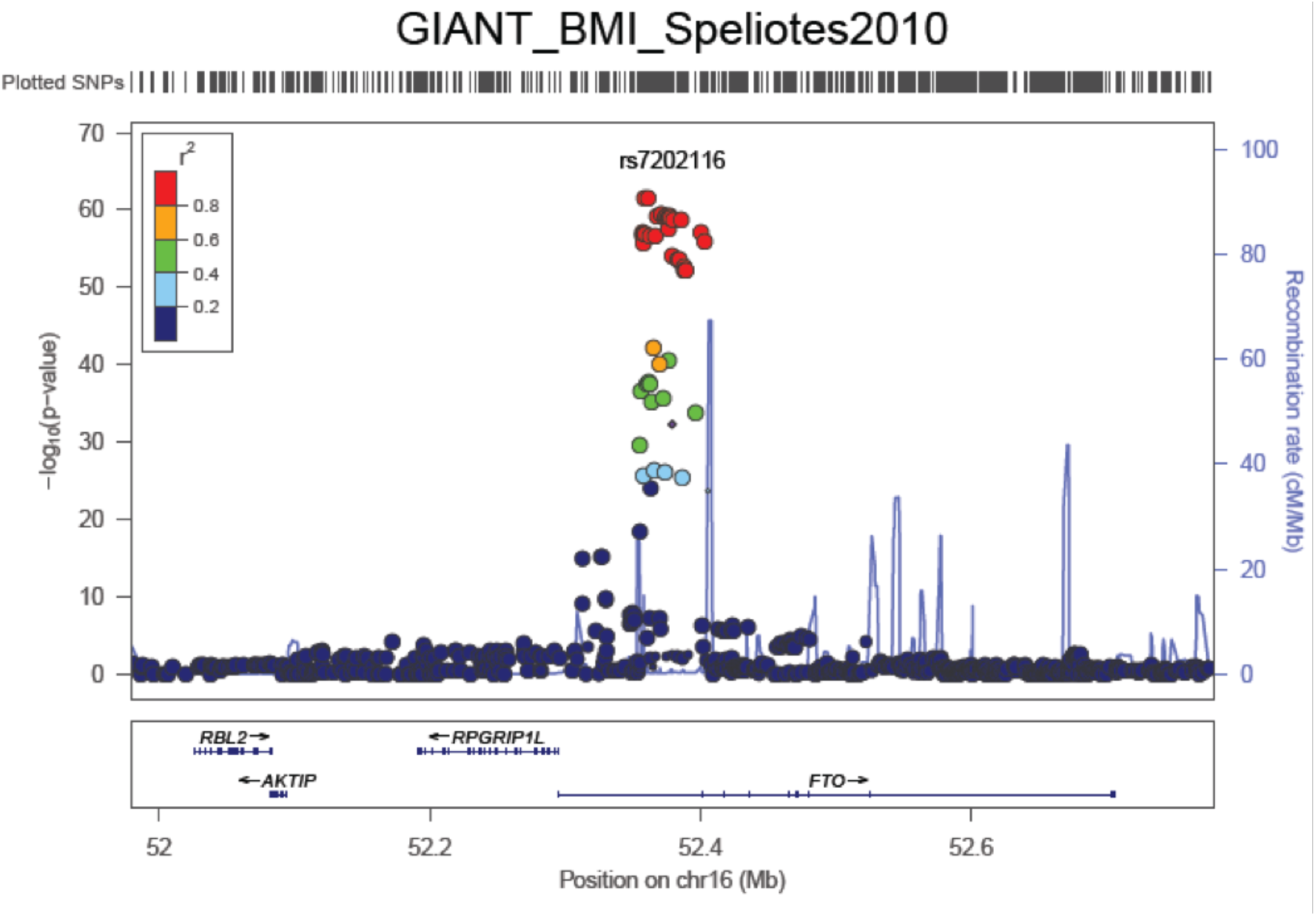
Regional Association Plot of rs7202116, top hit for variance in BMI found by Yang et al. (2012), on mean level of BMI from GIANT consortium data. Figure produced using LocusZoom ( [60]).

**S7 Fig.**
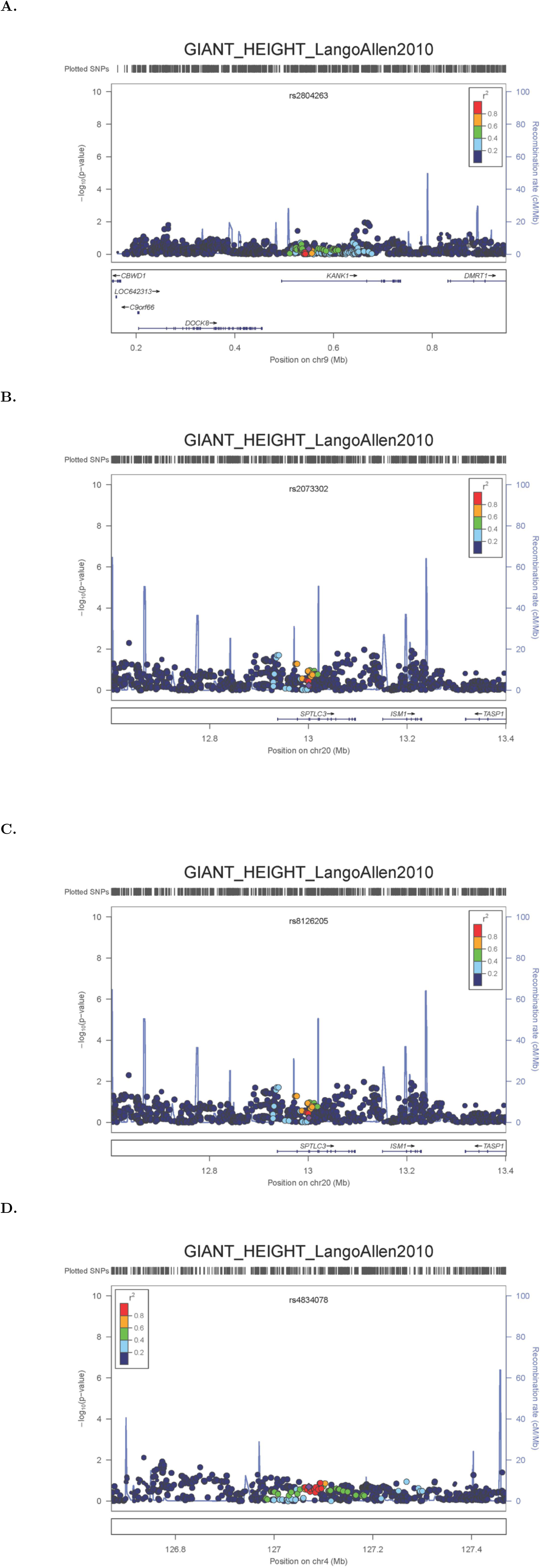
Regional Association Plot of genome-wide suggestively significant (p<10−5) hits from Fig 1 on mean height from GIANT consortium data. (A-D) Plots for the SNPs rs2804263, rs2073302, rs8126205, and rs4834078, respectively, show no markers in the respective regions that approach even genome-wide suggestive significance (p< 10^−5^). Figures produced using LocusZoom ( [60]).

**S8 Fig.**
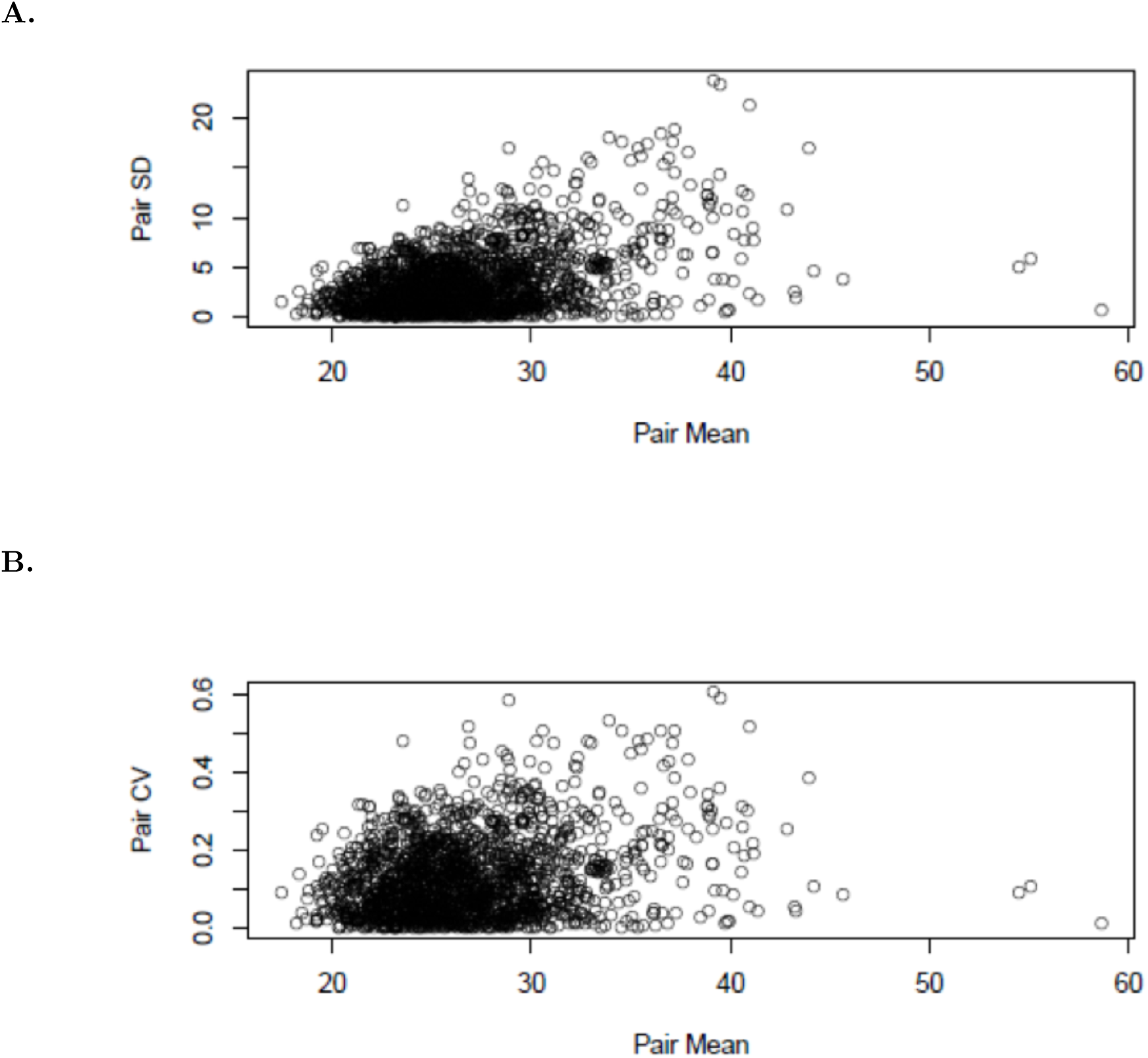
Relationship between sibling-pair mean BMI and sibling-pair standard deviation (SD) or coefficient of variation (CV). A) Sibling-pair SD versus mean (*ρ* = 0.43). B) Sibling-pair CV versus mean (*ρ* = 0.25).

**S9 Fig.**
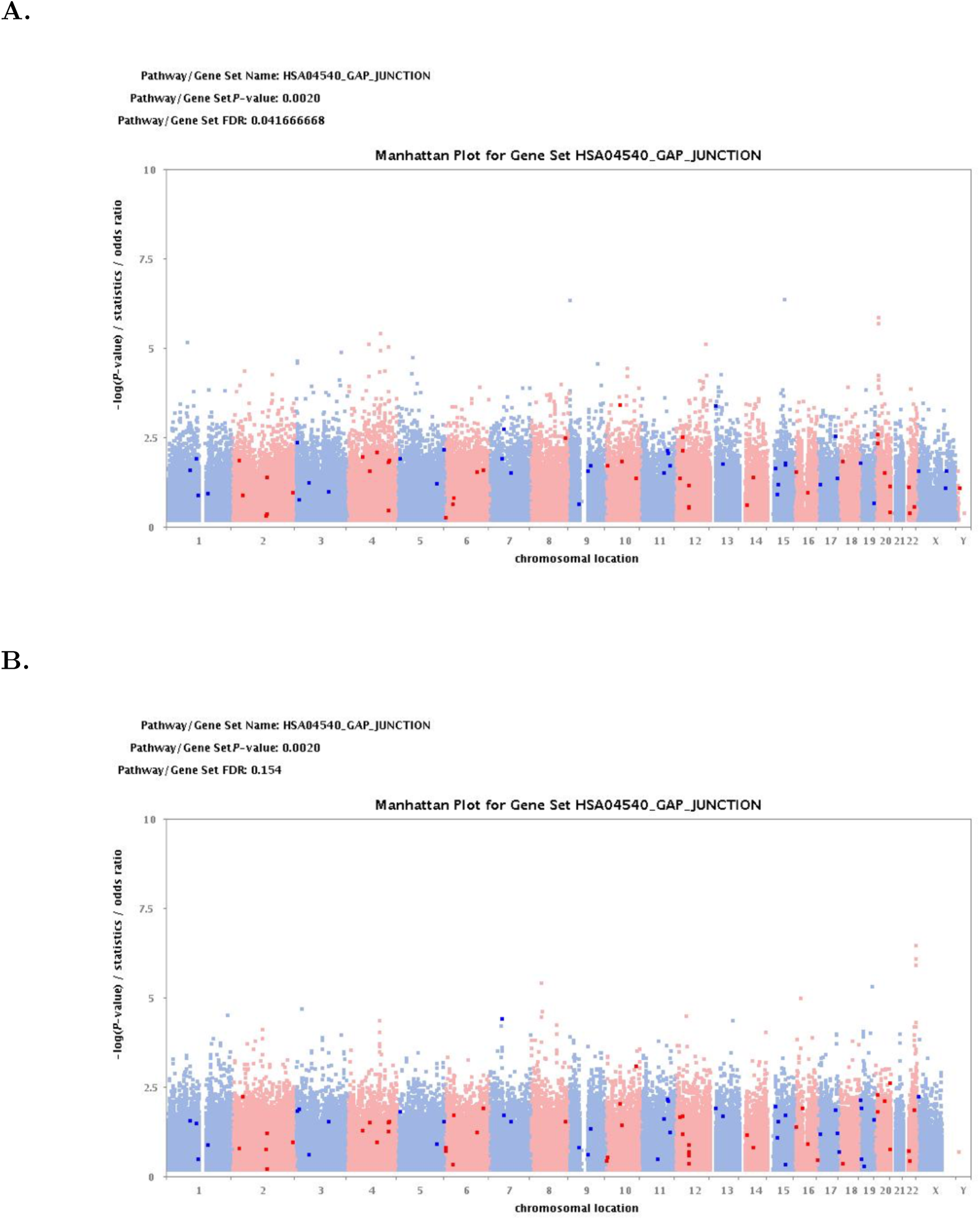
Manhattan plots for enriched pathway HSA04540 Gap Junction for height variability. A) FHS discovery sample; B) MTFS replication sample.

**S1 Table.** Pooled regression model of trait with no mean or variance effects on minor allele count with controls for sex and age. The table summarizes results for one randomly chosen replicate. For each of the models, one sibling out of each pair was randomly drawn. Controls for population stratification were included via an indicator variable for the one of the four subpopulations that generated the parents’ and offspring’s genotype. The results show that while the data with no confounding between genotype and the outcome variable correctly fails to reject the null of no effect, the pooled regression returns upwardly biased results in the presence of family-level confounding. Controls for broad population stratification do not successfully reduce this bias.

| | High degree of<br>family-level<br>confounding<br>($\rho = 0.3$) | | Medium degree of<br>family-level<br>confounding<br>($\rho = 0.15$) | | No<br>family-level<br>confounding<br>($\rho = 0$) | |
| --- | --- | --- | --- | --- | --- | --- |
| | $\beta$<br>($p$) | $\beta$<br>($p$) | $\beta$<br>($p$) | $\beta$<br>($p$) | $\beta$<br>($p$) | $\beta$<br>($p$) |
| Individual<br>minor<br>allele count | 0.4839<br>($< 0.0001$) | 0.4842<br>($< 0.0001$) | 0.2054<br>($< 0.0001$) | 0.2033<br>($< 0.0001$) | -0.0131<br>(0.7295) | -0.0118<br>(0.7569) |
| Controls for<br>pop strat | No | Yes | No | Yes | No | Yes |
| Observations | 4000 | 4000 | 4000 | 4000 | 4000 | 4000 |

**S2 Table.** Random effects regression model of trait with no mean or variance effects on minor allele count with controls for sex and age. The table summarizes results for one randomly chosen replicate. Random effects regressions were fit using the “random” option in R’s *plm* package using the default estimation method. The sample size is *N* = 8000 rather than *N* = 4000 because both offspring in a family unit are used. The results show that the estimates for the coefficients are less biased than in the pooled model (shown in greater detail across replicates (see Results)) but that in the presence of non-zero confounding between genotype and outcome, there is upward bias in the coefficients.

| | High degree of<br>family-level<br>confounding<br>($\rho = 0.3$) | | Medium degree of<br>family-level<br>confounding<br>($\rho = 0.15$) | | No<br>family-level<br>confounding<br>($\rho = 0$) | |
| --- | --- | --- | --- | --- | --- | --- |
| | $\beta$<br>($p$) | $\beta$<br>($p$) | $\beta$<br>($p$) | $\beta$<br>($p$) | $\beta$<br>($p$) | $\beta$<br>($p$) |
| Individual<br>minor<br>allele count | 0.3229<br>($< 0.0001$) | 0.3239<br>($< 0.0001$) | 0.1397<br>($< 0.0001$) | 0.1400<br>($< 0.0001$) | 0.0085<br>(0.7493) | 0.0096<br>(0.7197) |
| Controls for<br>pop strat | No | Yes | No | Yes | No | Yes |
| Observations | 8000 | 8000 | 8000 | 8000 | 8000 | 8000 |

**S3 Table.** Fixed effects regression model of trait with no mean or variance effects on minor allele count with controls for sex and age. The table summarizes results for one randomly chosen replicate. The sample is *N* = 8000 because both offspring from a family were used and there are no controls for population stratification because the indicator for the subpopulation does not vary between siblings and thus drops out of the regression. The results show that across all three degrees of family-level confounding, the fixed effects regression correctly fails to reject the null of no effects of the minor allele count on the outcome.

| | High degree of<br>family-level<br>confounding<br>($\rho = 0.3$)<br>$\beta$<br>($p$) | Medium degree of<br>family-level<br>confounding<br>($\rho = 0.15$)<br>$\beta$<br>($p$) | No<br>family-level<br>confounding<br>($\rho = 0$)<br>$\beta$<br>($p$) |
| --- | --- | --- | --- |
| Individual<br>minor allele<br>count | 0.0351<br>(0.3567) | -0.0357<br>(0.2580) | 0.0351<br>(0.3567) |
| Observations | 8000 | 8000 | 8000 |

**S4 Table.** Results of Hausman test comparing 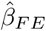 with 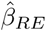 from S2 Table and S3 Table.

| Correlation level | % of simulations<br>that reject null<br>at $p < 0.05$ |
| --- | --- |
| 0 | 0.0570 |
| 0.15 | 0.9920 |
| 0.3 | 1.0000 |

**S5 Table.** Results of regressing squared Z-score of trait on minor allele count across 1000 replicates with non-demeaned data. The results show an inflated type I error rate for the trait simulated to have mean effects but no variance effects in the presence of an unobserved confounder between genotype and outcome (underlined rows). There is also a higher type I error rate than the sibling SD method for this trait even when there is no unobserved confounding.

| Simulated DV | Controls for ancestry? | Percent of sims with $p < 0.05$ on minor allele count |
| --- | --- | --- |
| *No confounding* |  |  |
| Neither mean nor var effects | no | 7.10 |
| Neither mean nor var effects | yes | 7.10 |
| Mean effects only | no | 7.10 |
| Mean effects only | yes | 6.50 |
| Var effects only | no | 100 |
| Var effects only | yes | 100 |
| Mean + var effects | no | 100 |
| Mean + var effects | yes | 100 |
| *Some confounding ($\rho = 0.15$)* |  |  |
| Neither mean nor var effects | no | 8.00 |
| Neither mean nor var effects | yes | 7.90 |
| <u>Mean effects only</u> | no | 24.60 |
| <u>Mean effects only</u> | yes | 24.00 |
| Var effects only | no | 100 |
| Var effects only | yes | 100 |
| Mean + var effects | no | 100 |
| Mean + var effects | yes | 100 |

**S6 Table.** Results of regressing squared Z-score of trait on minor allele count across 1000 replicates. Regressions are estimated using the demeaned data. The results show an inflated type I error rate for the trait simulated to have mean effects but no variance effects in the presence of an unobserved confounder between genotype and outcome (underlined row) even after transforming the data.

| Simulated DV | Percent of sims with $p < 0.05$<br>on minor allele count |
| --- | --- |
| *Some confounding* ($\rho = 0.15$) | |
| Neither mean nor var effects | 10 |
| <u>Mean effects only</u> | 20 |
| Var effects only | 90 |
| Mean + var effects | 100 |

**S7 Table.** Results of DGLM on non-demeaned (non-transformed) and demeaned data (transformed) for simulated DV with *variance effects only* across 1000 replicates. The results show an inflated type I error rate (estimate *β* ≠ 0 despite the presence of allele affects on the variance and not the mean) that is smaller but still present in the demeaned data. The results also show that while demeaning reduces the type I error rate (false detection of mean effects), the transformation leads to type II errors (fails to detect variance effects when these are present).

| Coefficient on minor allele count | Type of data | % coef $\neq 0$ at $p < 0.05$ |
| --- | --- | --- |
| *No confounding* |  |  |
| $\beta$ (mean; false positive) | Demeaned | 19.9 |
| $\beta$ (mean; false positive) | Non-transformed | 13.3 |
| $\gamma$ (variance; true positive) | Demeaned | 0 |
| $\gamma$ (variance; true positive) | Non-transformed | 100 |
| *Some confounding* ($\rho = 0.15$) | | |
| $\beta$ (mean; false positive) | Demeaned | 20.4 |
| $\beta$ (mean; false positive) | Non-transformed | 100 |
| $\gamma$ (variance; true positive) | Demeaned | 0 |
| $\gamma$ (variance; true positive) | Non-transformed | 99.9 |

**S8 Table.** Regression of sibling standard deviation in a trait on sibling count of minor alleles: one randomly chosen replicate. *Does not* control for parental genotype but controls for: sex of each offspring, age of each offspring; ancestry indicator. The results show that the method detects variance effects when these are present in the simulated dependent variable and correctly rejects the minor allele count leading to an increase in the sibling standard deviation for the dependent variable simulated to have mean effects only.

|  | *Simulated dependent variable:* |  |  |  |  |  |  |  |
| --- | --- | --- | --- | --- | --- | --- | --- | --- |
| | No family-level confounder ($\rho = 0$) | | | | Family-level confounder ($\rho = 0.15$) | | | |
|  | No effect | Mean effect | Var effect | Var and<br>mean effect | No effect | Mean effect | Var effect | Var and<br>mean effect |
| Sibling<br>minor<br>allele<br>count | 0.012<br>(0.011) | −0.003<br>(0.011) | 0.094***<br>(0.013) | 0.106***<br>(0.013) | 0.013<br>(0.009) | 0.011<br>(0.009) | 0.082***<br>(0.010) | 0.072***<br>(0.010) |
| Sibling<br>mean of<br>trait | 0.002<br>(0.009) | −0.014<br>(0.009) | −0.008<br>(0.010) | 0.006<br>(0.010) | −0.003<br>(0.009) | −0.032***<br>(0.009) | −0.003<br>(0.010) | 0.013<br>(0.010) |
| Observations | 4,000 | 4,000 | 4,000 | 4,000 | 4,000 | 4,000 | 4,000 | 4,000 |
| Adjusted R<sup>2</sup> | 0.001 | −0.001 | 0.013 | 0.016 | 0.005 | 0.003 | 0.015 | 0.013 |
| F Statistic<br>(df = 6; 3993) | 1.435 | 0.557 | 9.875*** | 11.616*** | 4.536*** | 3.129*** | 10.966*** | 10.054*** |

*Note:*
\*p<0.1; \*\*p<0.05; \*\*\*p<0.01

**S9 Table.** Regression of sibling standard deviation in a trait on sibling count of minor alleles: one randomly chosen replicate. *Does* control for parental genotype, as well as sex of each offspring, age of each offspring; ancestry indicator. The results show that the method detects variance effects when these are present in the simulated dependent variable and correctly rejects the minor allele count leading to an increase in the sibling standard deviation for the dependent variable simulated to have mean effects only.

|  | *Simulated dependent variable:* |  |  |  |  |  |  |  |
| --- | --- | --- | --- | --- | --- | --- | --- | --- |
| | No family-level confounder ($\rho = 0$) | | | | Family-level confounder ($\rho = 0.15$) | | | |
|  | No effect | Mean effect | Var effect | Var and<br>mean effect | No effect | Mean effect | Var effect | Var and<br>mean effect |
| Sibling<br>minor<br>allele<br>count | 0.022<br>(0.018) | 0.016<br>(0.018) | 0.102***<br>(0.022) | 0.091***<br>(0.022) | 0.014<br>(0.016) | 0.009<br>(0.016) | 0.100***<br>(0.018) | 0.054***<br>(0.018) |
| Sibling<br>mean of<br>trait | 0.002<br>(0.009) | -0.013<br>(0.009) | -0.008<br>(0.010) | 0.006<br>(0.010) | -0.003<br>(0.009) | -0.032***<br>(0.009) | -0.003<br>(0.010) | 0.013<br>(0.010) |
| Observations | 4,000 | 4,000 | 4,000 | 4,000 | 4,000 | 4,000 | 4,000 | 4,000 |
| Adjusted R<sup>2</sup> | 0.001 | -0.001 | 0.013 | 0.016 | 0.005 | 0.003 | 0.015 | 0.014 |
| F Statistic<br>(df = 7; 3992) | 1.296 | 0.707 | 8.492*** | 10.050*** | 3.888*** | 2.684*** | 9.626*** | 8.835*** |

*Note:*
\*p<0.1; \*\*p<0.05; \*\*\*p<0.01

**S10 Table.** Results of regressing sibling SD of trait on minor allele count across 1000 replicates. The results show that in contrast to the squared Z-score and DGLM, which each, in the presence of an unobserved confounder, display type I error rates of around 20% in detecting variance effects in traits simulated to have mean effects only, the sibling SD method avoids this type of error (underlined rows) both with and without controls for parental genotype. The results also illustrate that the method detects variance effects when the trait either has variance effects only or when the trait exhibits both mean and variance effects. The first half of the table also shows a lower type I error rate than squared Z-score when there is no unobserved confounder.

| Simulated DV | Family-level<br>confounding $\rho = 0.15$ | Controls for<br>parent genotype? | Percent of sims with $p < 0.05$<br>on minor allele count |
| --- | --- | --- | --- |
| *No confounding* |  |  |  |
| Neither mean nor var effects | No | No | 5.00 |
| Neither mean nor var effects | No | Yes | 4.80 |
| Mean effects only | No | No | 5.60 |
| Mean effects only | No | Yes | 4.80 |
| Variance effects only | No | No | 100.00 |
| Variance effects only | No | Yes | 92.40 |
| Mean + var effects | No | No | 99.70 |
| Mean + var effects | No | Yes | 92.00 |
| *Some confounding ($\rho = 0.15$)* |  |  |  |
| Neither mean nor var effects | Yes | No | 4.90 |
| Neither mean nor var effects | Yes | Yes | 5.80 |
| Mean effects only | Yes | No | 5.20 |
| Mean effects only | Yes | Yes | 4.80 |
| Variance effects only | Yes | No | 99.70 |
| Variance effects only | Yes | Yes | 91.60 |
| Mean + var effects | Yes | No | 99.70 |
| Mean + var effects | Yes | Yes | 90.70 |

**S11 Table.** Proxy SNPs and results for replication analysis using Minnesota Twin Family Study data.

| **FHS SNP** | **MTFS Proxy** | **Distance** | **$R^2$** | **D** | **P-value** | **N (pairs)** |
| --- | --- | --- | --- | --- | --- | --- |
| **Height SNPs** |  |  |  |  |  |  |
| rs2804263 | rs2804279 | 8976 | 0.959 | 1 | 0.174 | 1555 |
| rs2073302 | N/A | – | – | – | – |  |
| rs8126205 | N/A | – | – | – | – |  |
| rs4834078 | rs4348108 | 14075 | 1 | 1 | 0.782 | 1555 |
| rs4834078 | rs11730519 | 23237 | 0.959 | 1 | 0.89 | 1555 |
| rs4834078 | rs6852484 | 3263 | 0.851 | 1 | 0.75 | 1555 |
| **BMI SNPs** |  |  |  |  |  |  |
| rs30731 | rs28636 | 510 | 0.85 | 1 | 0.028 | 1555 |
| rs41508049 | rs10261193 | 1624 | 0.84 | 1 | 0.204 | 1555 |

**S12 Table.** Replicated GWAS hits for other SNPs on MAST4. Results are from the NCBI Phenotype-Genotype Integrator

| Trait | SNP | P.Value |
| --- | --- | --- |
| Autistic Disorder | 29456 | 0.00000 |
| Autistic Disorder | 17197559 | 0.00003 |
| Autistic Disorder | 1864036 | 0.00003 |
| Autistic Disorder | 253234 | 0.00004 |
| Blood Pressure | 10515002 | 0.00000 |
| Blood Pressure | 1366275 | 0.00001 |
| Body Height | 1363935 | 0.00005 |
| Body Height | 1363933 | 0.00006 |
| Body Height | 10500564 | 0.00006 |
| Body Mass Index | 25832 | 0.00000 |
| Body Weights and Measures | 1363935 | 0.00006 |
| Bronchodilator Agents | 146002062 | 0.00000 |
| Bronchodilator Agents | 189845032 | 0.00000 |
| C-Reactive Protein | 10056426 | 0.00006 |
| C-Reactive Protein | 26923 | 0.00009 |
| C-Reactive Protein | 26929 | 0.00010 |
| Carotid Intimal Medial Thickness 1 | 1697137 | 0.00000 |
| Child Development Disorders, Pervasive | 2801640 | 0.00001 |
| Child Development Disorders, Pervasive | 2968192 | 0.00001 |
| Child Development Disorders, Pervasive | 2561078 | 0.00003 |
| Chlorine | 6878808 | 0.00001 |
| Cholesterol | 1364020 | 0.00000 |
| Cholesterol, LDL | 1364020 | 0.00002 |
| Diabetes Mellitus, Type 2 | 1030231 | 0.00009 |
| Electrocardiography | 10514995 | 0.00000 |
| Epilepsy | 39861 | 0.00000 |
| Forced Expiratory Volume | 189845032 | 0.00000 |
| gamma-Linolenic Acid | 1007500 | 0.00001 |
| HIV-1 | 1697137 | 0.00000 |
| Intercellular Adhesion Molecule-1 | 7714441 | 0.00000 |
| Linoleic Acid | 17278159 | 0.00000 |
| Linoleoyl-CoA Desaturase | 1007500 | 0.00001 |
| Lipids | 1469419 | 0.00001 |
| Lung Volume Measurements | 146002062 | 0.00000 |
| Lymphocyte Count | 16895456 | 0.00005 |
| Lymphocyte Count | 12055346 | 0.00008 |
| Personality | 10052424 | 0.00006 |
| Platelet Function Tests | 9687339 | 0.00000 |
| Platelet Function Tests | 16895178 | 0.00000 |
| Platelet Function Tests | 2545386 | 0.00000 |
| Platelet Function Tests | 10940079 | 0.00001 |
| Platelet Function Tests | 4700148 | 0.00001 |
| Platelet Function Tests | 3111632 | 0.00001 |
| Platelet Function Tests | 10057708 | 0.00001 |
| Resistin | 253234 | 0.00001 |
| Resistin | 29456 | 0.00006 |
| Sodium | 6878808 | 0.00002 |
| Triglycerides | 1469419 | 0.00001 |
| Triglycerides | 2561078 | 0.00004 |

**S13 Table.** Gene set analysis results using PASCAL The table shows significant gene sets in FHS that replicated in MTFS at different p-value thresholds (MAST4, the location of the replicated SNP, does not appear because although it was *p* < 0.01 in the FHS dataset, it was *p* = 0.1 in the MTFS dataset).

| P-value threshold | Gene sets significant in FHS and MTFs |
| --- | --- |
| ***Height*** |  |
| 0.050 | CMPK1,FOXE3,FOXD2-AS1,FOXD2,LOC441601,MIR4692,EEF1DP3,SCG5,SPECC1,CCDC144CP,DLGAP1-AS5,C19orf40,NANP,LOC100134868,LOC284801,MIR26A1,PARL,LARP1B,FBXO8,LOC100506548,RPL37,SNORD72,CARD6,STAG3L4,EN2,CNPY1,FAM219A,DNAI1 |
| 0.01 | EN2 |
| ***BMI*** |  |
| 0.05 | NOL9,TAS1R1,NAV1,MIR5191,DUPD1,MPPED2,TCHP,FBF1,KIRREL2,APLP1,LRFN1,GMFG,SAMD4B,RELB,CLASRP,ZNF296,GEMIN7,PPP1R37,TTC27,GPR55,SENP2,CPEB2,CDC20B,MIR449C,UST,MAD1L1 |
| 0.01 | CLASRP,ZNF296,GEMIN7 |

**S14 Table.** Pathway analysis results using PASCAL The table shows significant pathways in FHS that replicated in MTFS at different p-value thresholds. The pathway that replicated using the **i-GSEA4GWAS** tool is not among those tested by **PASCAL**

| P-value threshold | Pathways significant in FHS and MTFs |
| --- | --- |
| ***Height*** |  |
|  | None |
| ***BMI*** |  |
| 0.05 | KEGG_ALPHA_LINOLENIC_ACID_METABOLISM<br>REACTOME_ACYL_CHAIN_REMODELLING_OF_PS<br>BIOCARTA_HER2_PATHWAY |

**S15 Table.** Illustrating correlation between population indicators and family-level intercept across 1000 replicates at three degrees of family-level confounding. The results show that at higher degrees of confounding between an unobserved characteristic of a family and observed genotype, there is a stronger relationship between the indicator for the respondent’s ancestry and the family-level intercept. This shows that a control for ancestry can attenuate some bias in the estimate created by confounding but not fully eliminate this bias.

| $cor(X_{ij}, \alpha_j)$ | Percent significant<br>$\beta$ on<br>population<br>indicator |
| --- | --- |
| 0 | 15.9% |
| 0.15 | 28.9% |
| 0.30 | 48.6% |

**S16 Table.** Relationship between family intercept and observed genotype. The table shows that as the degree of between-family confounding increases, there is a stronger relationship between the intercept that shifts levels of a trait up or down between families and the genotype.

| $cor(X_{ij}, \alpha_j)$ | Mean $\beta_1$ |
| --- | --- |
| 0 | 0.0003 |
| 0.15 | 0.1766 |
| 0.30 | 0.3534 |

